# Merging Metabolic Modeling and Imaging for Screening Therapeutic Targets in Colorectal Cancer

**DOI:** 10.1101/2024.05.24.595756

**Authors:** Niki Tavakoli, Emma J. Fong, Abigail Coleman, Yu-Kai Huang, Mathias Bigger, Michael E. Doche, Seungil Kim, Heinz-Josef Lenz, Nicholas A. Graham, Paul Macklin, Stacey D. Finley, Shannon M. Mumenthaler

## Abstract

Cancer-associated fibroblasts (CAFs) play a key role in metabolic reprogramming and are well-established contributors to drug resistance in colorectal cancer (CRC). To exploit this metabolic crosstalk, we integrated a systems biology approach that identified key metabolic targets in a data-driven method and validated them experimentally. This process involved a novel machine learning-based method to computationally screen, in a high-throughput manner, the effects of enzyme perturbations predicted by a computational model of CRC metabolism. This approach reveals the network-wide effects of metabolic perturbations. Our results highlighted hexokinase (HK) as a crucial target, which subsequently became our focus for experimental validation using patient-derived tumor organoids (PDTOs). Through metabolic imaging and viability assays, we found that PDTOs cultured in CAF-conditioned media exhibited increased sensitivity to HK inhibition, confirming the model predictions. Our approach emphasizes the critical role of integrating computational and experimental techniques in exploring and exploiting CRC-CAF crosstalk.

## Introduction

Tumor metabolism, recognized as a hallmark of cancer, is increasingly garnering attention as an avenue for therapeutic intervention.^1–3^ A number of metabolic vulnerabilities have been identified; however, to date, efforts to bring metabolism-based drugs to the clinic have shown limited success. A significant hurdle lies in the intricate metabolic interplay within tumors, particularly the interactions between cancerous cells and nonmalignant cells within the tumor microenvironment (TME).^4^ For example, studies have demonstrated a co-survival relationship between tumor cells and cancer associated fibroblast (CAFs), highlighting reciprocal interactions involving nutrient secretion and consumption.^5^ This metabolic reprogramming has been shown to support tumor growth and drug resistance.^6–8^ Given this complexity, utilizing a systems biology approach that combines high-throughput computational and experimental techniques is beneficial, as recently reviewed in Curvello *et al*.^9^

In the context of tumor metabolism, modeling has produced relevant quantitative insights in previous work.^10^ Metabolic modeling encompasses various different techniques, each with its unique applications and insights.^11^ One such technique is kinetic modeling, which simulates the dynamics of a system, predicting how the metabolite concentrations and reaction rates evolve over time. For example, in pancreatic cancer research, kinetic models have been fitted to experimental data to gain deeper insights into the network dynamics and perform enzyme knockdowns to identify ways to inhibit cancer cell proliferation.^12^ Agent-based modeling is a technique that represents individual cells interacting with the TME and has been used to show metabolic changes under different environments.^13^ Another technique is multi-scale modeling, which spans various spatial and temporal scales, offering a comprehensive view of biological processes. Such models incorporate intracellular models of metabolism to inform cell- and tissue-level behaviors.^14,15^ Lastly, constraint-based modeling stands out as a widely adopted form of metabolic modeling. This technique works by predicting the flux distributions within a network by applying a series of constraints, allowing for the exploration of metabolic behaviors under different physiological and pathological conditions. Notably, constraint-based models can incorporate multi-omics data to generate context-specific models.^16–18^ Additionally, performing flux balance analysis^19^ using a constraint-based model of metabolism can predict enzymes that are the most essential to target and exploit.^20,21^

Our previous study focused on mechanisms underlying the metabolic relationship between colorectal cancer (CRC) cells and CAFs.^22^ By integrating metabolomics data and predefined experimental growth rates, constraint-based modeling revealed distinct ways in which CAFs specifically reprogram the central carbon metabolism of CRC cells. Notably, the presence of CAFs resulted in a significant upregulation of glycolysis, inhibition of the tricarboxylic acid (TCA) cycle and a disconnection in the oxidative and non-oxidative arms of the pentose phosphate pathway (PPP). The study also found distinct CAF-mediated alterations in glutamine metabolism. In addition, we applied the model to identify metabolic perturbations that uniquely inhibit growth of CRC cells in CAF-conditioned media (CAF-CM), compared to their original media. While informative, this prior work did not evaluate network-wide effects of metabolic perturbations. Furthermore, we did not experimentally validate the predicted targets.

In this current study, we significantly move beyond the prior work to exploit the metabolic relationship between CRC cells and CAFs. We developed a novel systems biology workflow (**Figure 1**) for high throughput computational screening aimed at identifying metabolic perturbations that could serve as potential druggable targets. Using representation learning to reduce the dimensionality of our data^23^, we performed a series of *in silico* enzyme knockdowns and analyzed their effects on each reaction in the metabolic network. Among the identified potential targets, hexokinase (HK), one of the most impactful, became our perturbation of interest to experimentally validate. This experimental validation was performed using 3D patient-derived tumor organoids (PDTOs), a physiologically relevant cell culture model system that has been shown to recapitulate the genetic and phenotypic properties of the original tumor.^24–26^ We assessed drug responses using conventional cell viability assays and metabolic imaging techniques, namely fluorescence lifetime imaging microscopy (FLIM)^27,28^, to validate HK as a promising druggable target within the tumor microenvironment (TME). This refined focus underscores the unique metabolic dependencies created by CAF-conditioned media and emphasizes the need for our novel approach in uncovering the most promising therapeutic interventions for CRC in the TME.^29^

**Figure 1:**
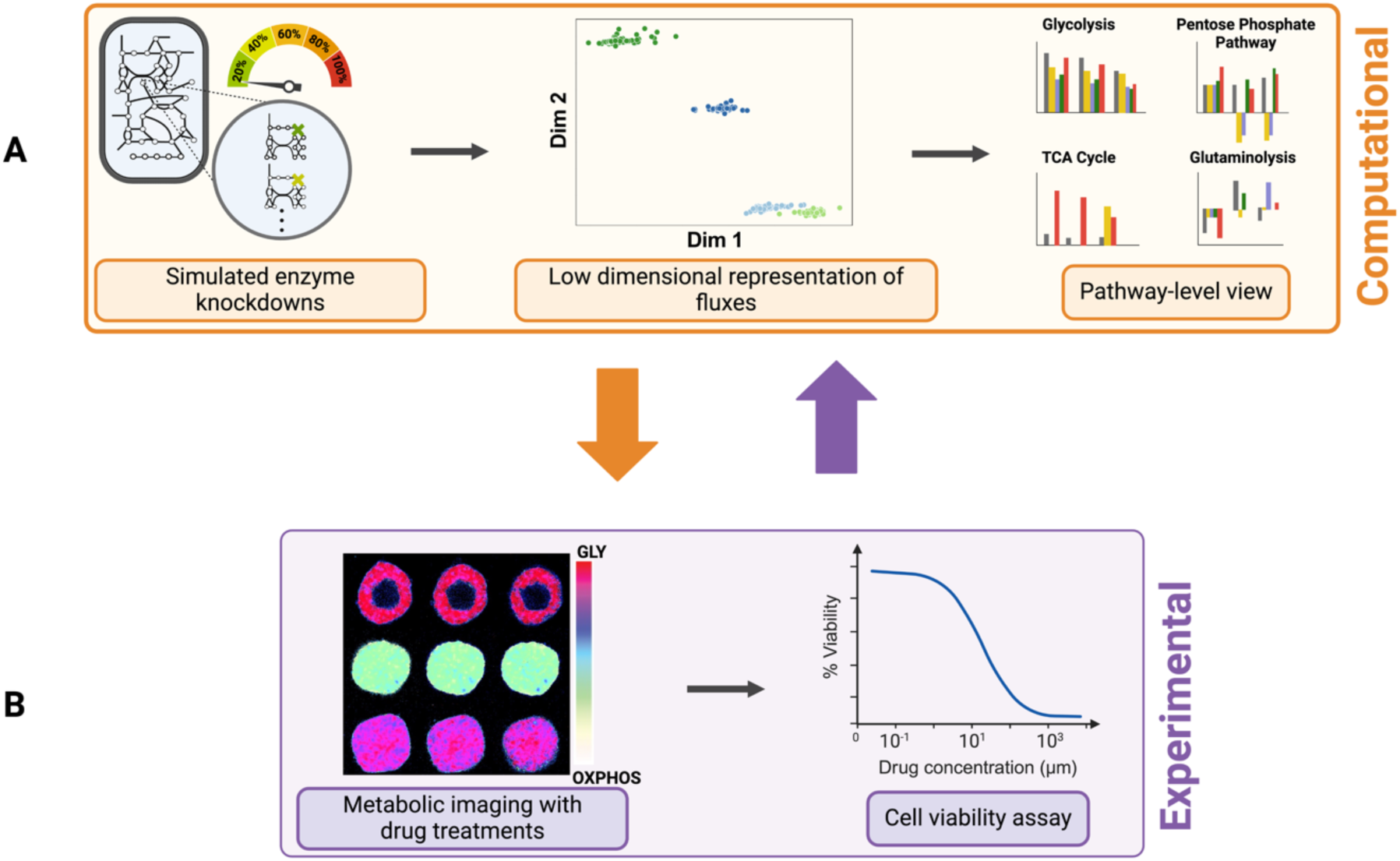
Project pipeline. **(A)** Computational portion of the study. Left: for each culture condition, metabolic reactions were subjected to knockdowns from zero (baseline) to 100% (full knockout) in increments of 20%. Middle: we applied a representation learning approach to reduce the dimensionality of our generated data, which enabled identification of enzyme perturbations that strongly impact the metabolic network. Right: to explore their effects, these flux changes were compared against baseline conditions (no knockdown), as well as against non-significant perturbations across the same and distinct cell conditions. **(B)** Experimental validation of *in silico* results through measurement of metabolic signatures (e.g. FLIM; left) and drug response assays (right). We returned to the computational model to generate additional calculations to compare to the experimentally observed metabolic imaging and drug responses, symbolizing the bidirectional arrows between panels **A** and **B**.

## Results

### Constraint-based modeling of CRC central carbon metabolism predicts the distribution of fluxes for distinct growth changes

We used an existing model of the metabolic pathways in central carbon metabolism from Wang *et al.*^22^ to investigate the metabolic crosstalk between CRC cells and CAFs. As detailed in that study, unsteady-state parsimonious flux balance analysis is used to determine the flux for each reaction in the model of central carbon metabolism, given mass balance constraints and measured fold-changes from metabolomics data.^22^ We predicted flux through the metabolic network for four conditions using this method: KRAS mutant (KRAS^MUT^) and KRAS wildtype (KRAS^WT^) CRC cells cultured only in CRC media or switched to CAF-CM. For notation, we use WT_CRC_ and WT_CAF_ to denote KRAS^WT^ CRC cells cultured only in CRC media or switched to CAF-CM, respectively. Similarly, KRAS_CRC_ and KRAS_CAF_ refer to KRAS^MUT^ CRC cells cultured only in CRC media or switched to CAF-CM, respectively. Thus, we can compare the effects of the media or the KRAS mutation. We utilized the same metabolomics constraints and pre-defined growth rates as in our prior work, given in **Supplementary Table S1 and S2**.^22^ The predicted fluxes for KRAS^MUT^ cells considering the effect of the growth media are distinctly shown in **Supplementary Figure S1**. These results are the focus of this work, given the prevalence of colorectal cancers with the KRAS mutation and the overall worse prognosis; however, for completeness, predictions to investigate the effect of growth media and gene mutation are shown in **Supplementary Figures S2 – S4.** For the remainder of this work, we focus on KRAS^MUT^ cells in the two media types. The predicted flux distributions for KRAS^MUT^ cells were then used as an initial point for performing a wide set of enzyme knockdowns in the network and serve as the “baseline” points in our study, where there is no perturbation.

### Heatmap visualization of enzyme knockdowns in central carbon metabolism unveils widespread effects and dynamic responses

In addition to understanding the predicted flux distributions for the various conditions, we simulated perturbations to investigate the network-wide responses to enzyme knockouts. We aimed to expand the efforts of Wang *et al*^22^, whose work concentrated solely on the effects of complete enzyme knockouts on the biomass reaction. It is common to study how biomass is affected by metabolic perturbations; however, this approach does not produce a broader understanding of the network-level response to alterations. Therefore, we fully inhibited the activity of each enzyme within the network and looked at its resultant impact on overall network functionality. We further developed this analysis to include partial enzyme knockdowns (at levels of 20%, 40%, 60%, and 80%) across the entire network.

We summarize and display the impact of 100% enzyme knockdowns for KRAS_CAF_ and KRAS_CRC_ specifically in the heatmap shown in **Figure 2**. Each row corresponds to a single enzyme knockdown in the network, while the columns correspond to the predicted fluxes of reactions for the given enzyme knockdown. Each box corresponds to a single flux value. The heatmap view allows us to examine flux changes after metabolic perturbations at the network level.

**Figure 2:**
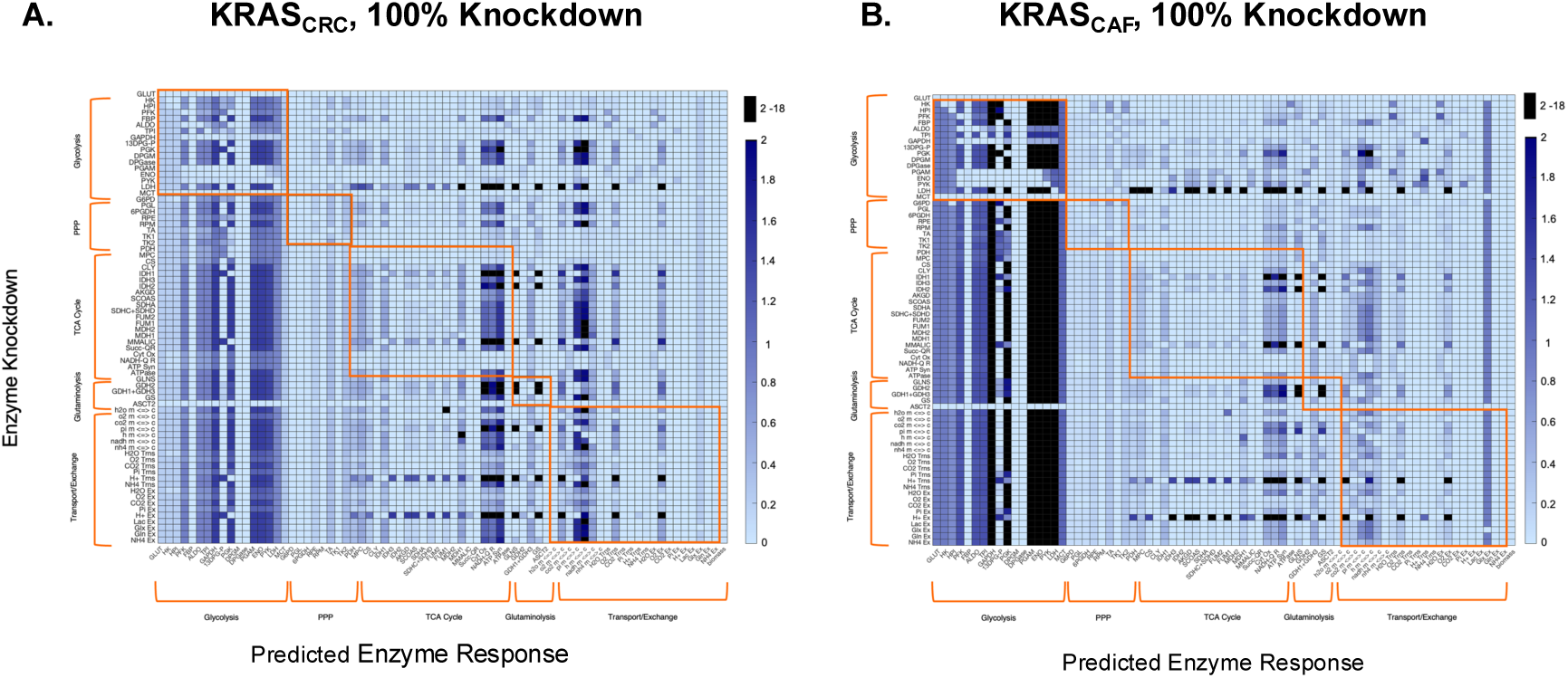
Heatmap displaying the impact of complete (100%) enzyme knockouts on absolute reaction fluxes. (A) KRAS_CAF_ condition and (B) KRAS_CRC_ condition. Rows represent individual enzyme knockdowns, while columns depict the effects of each perturbation on the rest of the network (predicted enzyme response). The color axis illustrates the absolute flux of enzymes after each perturbation, with color intensity indicating flux values from 0 to 2 hr^-1^, which covers the majority of the flux data. To enhance readability, any values exceeding 2 hr^-1^ have been masked as black. Outlined boxes highlight effects within individual pathways of central carbon metabolism, making it easier to visualize intra-pathway adjustments due to each knockdown.

In several instances, we see that the predicted fluxes are independent of which enzyme is inhibited, as indicated by the same colors extending vertically down the heatmap. For example, the figure shows that, overall, metabolic perturbations in multiple pathways lead to higher glycolytic flux, as indicated by the darker blue hue extending vertically within the block of glycolysis reactions on the left side of the figure. Additionally, following knockdown of most enzymes in the network, certain enzymes such as enolase (ENO), pyruvate kinase (PYK), and lactate dehydrogenase (LDH) are predicted to exhibit high flux values, indicated by the dark hue extending vertically. Therefore, enzymes not following suit are predicted to be causing a differential effect compared to other perturbations.

We also find that certain knockdowns cause unique effects within the network that are not seen in response to other metabolic perturbations. For instance, in glycolysis, the compared to other perturbations in the system. Further analysis indicates that these types of unique effects can be found within pathways other than just glycolysis. For example, knockout of the hydrogen ion exchange reaction also results in similar effects on TCA cycle fluxes, and this knockout also influences flux through the electron transport chain reactions. Contrarily, there may appear to be similarities in effects across these knockdowns; biologically, this could suggest a certain level of redundancy and robustness in the metabolic pathways. However, these differential effects are difficult to decipher efficiently by solely analyzing the heatmap. Therefore, we sought to develop an alternate strategy for identifying effective metabolic perturbations.

### Dimensionality reduction helps visualization and comparison of the effects of the metabolic perturbations

As denoted in **Figure 2**, heat maps can be used to visualize the network-level responses to inhibiting individual metabolic enzymes. However, it is difficult to compare across the conditions and perturbations, given the number of enzyme knockdowns simulated for each condition and the wide range of predicted flux values. Thus, we considered an alternative way to represent and visualize the predicted fluxes. We utilized a machine learning approach to perform dimensionality reduction and cast the distribution of fluxes within the network, considering enzyme knockdowns, into a 2D projection space (**Figure 3A**). This approach makes use of neural networks to reduce the dimensionality of model outputs and then to calculate the distance between the projected points to determine how different they are from each other.^30^ A detailed description of this approach is provided in the Methods section. Plotting the predicted reaction fluxes in 2D space is a vast reduction in the dimensionality of the model output, as each point represents the flux through all 74 reactions in response to a single enzyme knockdown. **Figure 3A** displays 1400 points (termed “coordinates”) in 2D space, derived from 4 conditions (WT_CRC_, WT_CAF_, KRAS_CRC_, and KRAS_CAF_), 70 enzyme knockdowns, and 5 different levels of inhibition. Each coordinate represents a single simulation, denoting a specific knockdown and its subsequent impact on the metabolic network, representing a row of the heatmap in **Figure 2**. The points denoted in red correspond to the calculated centroid (the “average” of all points for each condition).

**Figure 3:**
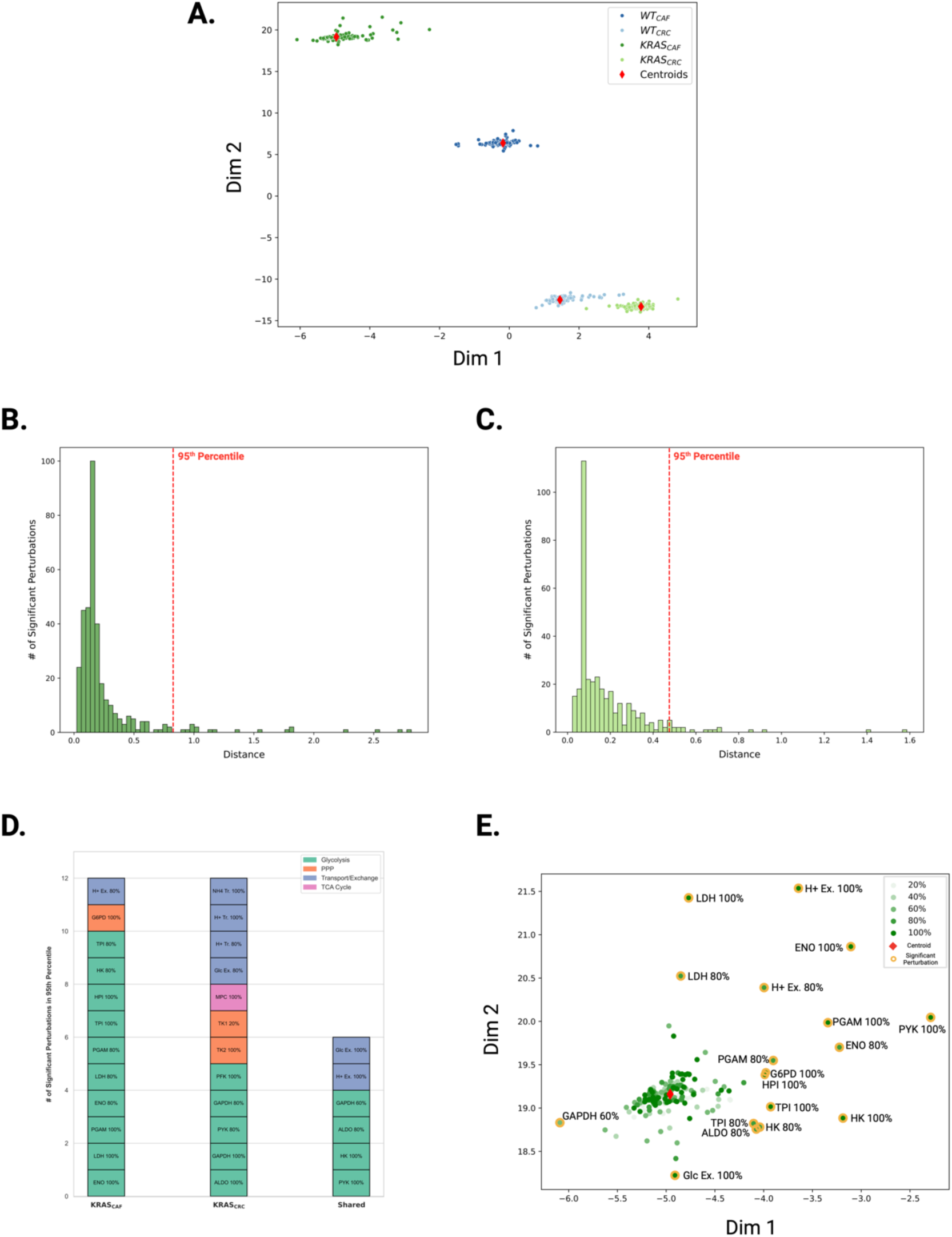
Dimensionality reduction and data analysis. **(A)** Predicted reaction fluxes due to metabolic perturbations projected in 2D space. **(B-C)** Distributions of distances between each simulation point in 2D space to its cluster centroid for KRAS_CAF_ and KRAS_CRC_ respectively. The dotted red lines represent the distance separating the top 5^th^ percentile of distances furthest from the centroid. **(D)** Influential metabolic perturbations identified for KRAS_CAF_ and KRAS_CRC_, according to central carbon metabolism pathways. **(E)** Labeling of simulated perturbations and identification of significant reactions in KRAS_CAF_ 2D projected space.

When the simulation coordinates are labeled according to their respective condition, they create distinct clusters, implying significant differences among the four conditions and reinforcing each condition’s uniqueness. We can infer the similarity of the predicted flux distributions based on how close the points are in the 2D projection space. While we cannot necessarily assign biological meaning to the two dimensions, based on the neural network training, the distance between points is a metric to represent their similarity (see Methods section for details). Thus, the grouping together of points in each condition indicates that they share similar characteristics. The two CAF-CM conditions (in darker colors) show more separation compared to the two CRC media conditions (in lighter colors). The coordinates corresponding to the KRAS_CAF_ condition are distinctly isolated from the rest of the simulation coordinates. We considered alternative dimensionality reduction methods, including principal component analysis (PCA), t-distributed stochastic neighbor embedding (t-SNE), uniform manifold approximation and projection (UMAP) (**Supplementary Figure S5**); however, these methods do not recover the distinct grouping of the four conditions simulated and cannot be used to compare the effects of the metabolic perturbations. We note that the t-SNE and UMAP utilize unsupervised learning methods, and cannot correctly cluster the complex, non-linear data generated by the gene knockdowns. In contrast, the dimensionality reduction method is a supervised learning approach using representation learning, in which the simulated gene knockdowns are fed into the neural network as labeled inputs representing four conditions. Thus, it is not surprising that our approach shows clustering of points within the four conditions. Our primary purpose of using representation learning is to identify metabolic perturbations that produce significantly altered flux distributions, rather than clustering, and we describe the steps to determine influential perturbations.

### Identification, classification, and analysis of influential perturbations for KRASCAF and KRASCRC

Next, we sought to identify significant metabolic perturbations of interest by examining the distributions of the Euclidean distances between each perturbed simulation coordinate and the calculated centroid for each condition. The distance between points in the projection space can be used to identify enzyme knockdowns that drastically shift the flux distribution from the base state (**Figure 3B-C** and **Supplementary Figure S6-9**). Our objective is to pinpoint perturbations that notably influence reaction flux values. We focus on the perturbations for KRAS^MUT^ cells, due to its clinical significance, and we consider the effect of CAF-CM due to the evidence in literature that CRC-CAF interactions promote drug resistance.^31,32^ We note that inputs to the SNN must have the same dimensions. As described in the Methods section, we trained the SNN to learn 2D representations of 70x74 matrices containing the predicted flux through 74 reactions when each of the 70 metabolic enzymes is perturbed. The baseline reaction flux vector is 1x74, and hence, cannot be inputted into the SNN since it is of a different dimension. Therefore, we consider the centroid of a cluster when identifying influential perturbations. Thus, we computed the Euclidean distances in the 2D learned space between the cluster centroid and each coordinate pertaining to KRAS_CAF_ and KRAS_CRC_ conditions and constructed separate histograms for each condition to study the distribution of these distances. For both conditions, the distances between the simulation coordinate and the centroid due to a metabolic perturbation fall between 0 and 0.25 (**Figure 3B-C**), indicating that the majority of perturbations produce network fluxes whose coordinate in the learned 2D space lie close to the centroid. Since all histograms presented as right-tailed distributions, we represented significant perturbations as the simulation coordinates that lie at the furthest end of the tail, which captures the most extreme deviations in each condition. Specifically, we use distances within the 95^th^ percentile for KRAS_CAF_ and KRAS_CRC,_ indicating the top 5% of coordinates with distances furthest from the centroid. Since each condition contained an equal number of points in the top 5th percentile (18 points), all of these influential perturbations were selected from each condition for a detailed comparative analysis, allowing us to effectively examine the impact of different growth conditions.

We classified the influential perturbations based on their metabolic pathway for both conditions, which revealed distinct differences for KRAS_CAF_ and KRAS_CRC_ (**Figure 3D**). KRASCAF contains significantly more glycolytic reactions, with them making up over 77% of the impact perturbations while glycolytic reactions only made up 50%. Importantly, the influential perturbations identified by our selection process for the KRAS_CAF_ condition include fully inhibited HK and glucose-6-phosphate dehydrogenase (G6PD), which were found to be reactions of importance in the work of Wang *et al*. as well.^22^Additionally, some reaction knockdowns are shown to be a significant perturbation in both conditions: HK, GAPDH, pyruvate kinase (PYK), aldolase (ALDO), hydrogen ion exchange, and glucose exchange. Thus, there are cases where an enzyme knockdown can alter network flux, regardless of the cell’s growth media. We can also identify metabolic perturbations that are unique to each condition; for example, G6PD 100% knockdown, a PPP enzyme, was other enzymes in PPP, TK1 and TK2, were found to uniquely affect the system in the KRAS_CRC_ condition. This highlights the differences in metabolic vulnerability and sensitivity caused by CAF-conditioned media.

We can also gain insights into how similarly the impactful metabolic perturbations are predicted to affect the flux distributions by considering their coordinates in the 2D projection space. To directly pinpoint where each of the significant perturbations fall in relation to one another, we labeled them on the scatterplot of 2D projections of enzyme knockdowns. We focus on the KRAS_CAF_ condition (**Figure 3E**). This analysis provides insight into the similarities in flux distributions in response to the metabolic perturbations predicted to be most influential. For instance, the coordinates for PYK 100%, phosphoglycerate mutase (PGAM) 100%, and ENO 80% are positioned closer together in one area of the projection space, whereas the 60% knockdown of glyceraldehyde-3-dehydrogenase (GAPDH) produces distinct metabolic effects that set them apart from the other impactful perturbations, on the opposite end of the plot. Furthermore, color-coding the points based on the strength of the knockdown indicates that the points farthest from the centroid do not exclusively correspond to complete enzyme inhibitions. Dimensionality reduction reveals that the total inhibition of a non-influential enzyme results in a position near the cluster centroid, which suggests that it closely resembles other perturbations with minimal system impact.

### The significant perturbations for KRASCAF produce diverse metabolic changes on the network using various mechanisms

The coordinates of the flux distributions given specific metabolic perturbations in 2D projection space, and their distances from the centroid, indicate the effect of a perturbation at the network level. We can further investigate the effects of metabolic perturbations on biomass production (a proxy for growth) and at the pathway level. We particularly focus on the KRAS_CAF_ condition and consider how flux through glycolysis, TCA cycle, the PPP, and glutaminolysis is altered.Our analysis of metabolic perturbations in the 2D projection space (**Figure 3**) identified the full knockout of PYK as the most significant perturbation. This was followed by the full knockouts of the hydrogen proton exchange reaction and ENO. Beyond these top-ranked perturbations in the KRAS_CAF_ condition, we observed that the majority of significant metabolic disruptions were associated with reactions in the intermediate and later stages of glycolysis. Among these, the knockdown of HK, which catalyzes the first and rate-limiting step of glycolysis, emerged as the 7^th^ most impact perturbation.

When selecting perturbations for experimental validation, we prioritized HK for several reasons. First, we recognize that proton exchange reactions between intracellular metabolites are not feasible to experimentally knock down and, therefore, lack biological relevance in the context of our model results. Second, numerous *in silico* studies have highlighted HK as a potentially vulnerable enzyme target for disrupting the metabolic crosstalk between CRC cells and CAFs.^10,33,34^ Additionally, since most significant perturbations in the KRAS_CAF_ condition were glycolytic in nature, we aimed to target an enzyme capable of effectively shutting down glycolysis entirely, preventing flux from intermediate knockdowns from propagating through the network. Given its role as the first and rate-limiting enzyme in glycolysis, HK was a logical choice. To further contextualize its impact, we also examined the effects of PYK knockout in the computational model. Such mechanistic understanding could be leveraged to improve clinical outcomes. We compared HK knockdown to both the baseline condition (with no perturbation) and to the knockdowns whose coordinates are in the left-most tail of the distribution of distances (bottom 5^th^ percentile). We consider those perturbations as “reference” points, as they are not predicted to significantly alter the baseline flux distribution due to their proximity to the centroid in the 2D projection space. We also considered the effect of completely inhibiting PYK, as well. This comparative analysis provided valuable insights into how different perturbations influence metabolic pathways within CRC cells.

We found that the knockouts of HK and PYK inhibit glycolysis and yield reduced fluxes, compared to both baseline and reference points, but in different ways. We found that the knockdown of HK resulted in an overall 29% reduction in flux across the pathway compared to baseline, with a minimal to slight increase in the LDH reaction (1.7%). PYK knockout resulted in an overall larger inhibition of flux through the pathway and reduced flux for the LDH reaction (1.203 hr^-1^ compared to 2.01 hr^-1^ in the baseline condition). Therefore, less lactate is produced and secreted from the cell in response to PYK shutdown. In the PPP, a distinct decoupling is observed between the oxidative and non-oxidative arms **(Figure 4B)**. The knockout of HK significantly reduces fluxes in the oxidative arm of the PPP, with a flux sum of 0.171 hr^-1^, compared to baseline and the mean of the reference points (0.257 hr^-1^ and 0.240 hr^-1^, respectively). It increases the flux in the non-oxidative arm, with a flux sum of 1.594 hr^-1^, compared to the baseline and mean of reference points (0.577 hr^-1^ and 0.664 hr^-1^, respectively). On the contrary, flux through PPP generally decreased with PYK knockdown. However, flux through G6PD and PGL in the non-oxidative arm were unchanged, and there was a notable increase in flux through the TK1 reaction in the oxidative arm (0.204 hr^-1^ compared to the baseline and mean of reference points 0.084 hr^-1^ and 0 hr^-1^, respectively). There were also substantial differences in fluxes in the TCA when comparing the HK and PYK knockdowns to the baseline and reference conditions (**Figure 4C**). The baseline and mean of reference points resulted in a flux sum of 0.867 hr^-1^ and 0.884 hr^-1^ respectively, while HK knockout resulted in a sum of 0.230 hr^-1^. This indicates a strong decrease of flux through the TCA cycle. In comparison, PYK knockout resulted in a flux sum of 2.845 hr^-1^, indicating a strong upregulation and buildup of TCA cycle intermediates. Lastly, the perturbations induced by HK and PYK knockouts result in alterations in glutaminolysis fluxes, as shown in **Figure 4D**. Specifically, the net flux through GDH increases from the baseline value of 0.046 hr⁻¹ to 0.050 hr⁻¹ for HK knockout and to 0.058 hr⁻¹ for PYK knockout. This shift reflects slightly enhanced utilization of glutamine, with a negative flux indicating the production of glutamate from α-ketoglutarate, while a positive flux signifies glutamate utilization for α-ketoglutarate synthesis. HK knockout notably upregulates flux through glutamine synthetase (GS), increasing from 0.0014 hr⁻¹ to 0.029 hr⁻¹, suggesting an upregulation of glutamine conversion to glutamate. In contrast, PYK knockout reduces the flux through GS, implying a diminished capacity for glutamine-to-glutamate conversion. Furthermore, the flux through glutaminase (GLS) is significantly elevated under both knockouts, with HK knockout increasing GLS flux to 0.220 hr⁻¹ and PYK knockout to 0.152 hr⁻¹. These findings indicate increased glutamine consumption and intracellular glutamine accumulation, particularly for HK knockout. In summary, we have systematically inhibited metabolic enzymes in a computational model of central carbon metabolism that accounts for metabolic crosstalk between CRC cells and CAFs. We evaluated how the metabolic perturbations affect flux through the network and identified which enzymes should be inhibited to significantly alter the network flux using a machine learning approach. This work provides a novel method of screening potential metabolic perturbations. We next sought to confirm our *in-silico* findings experimentally.

**Figure 4:**
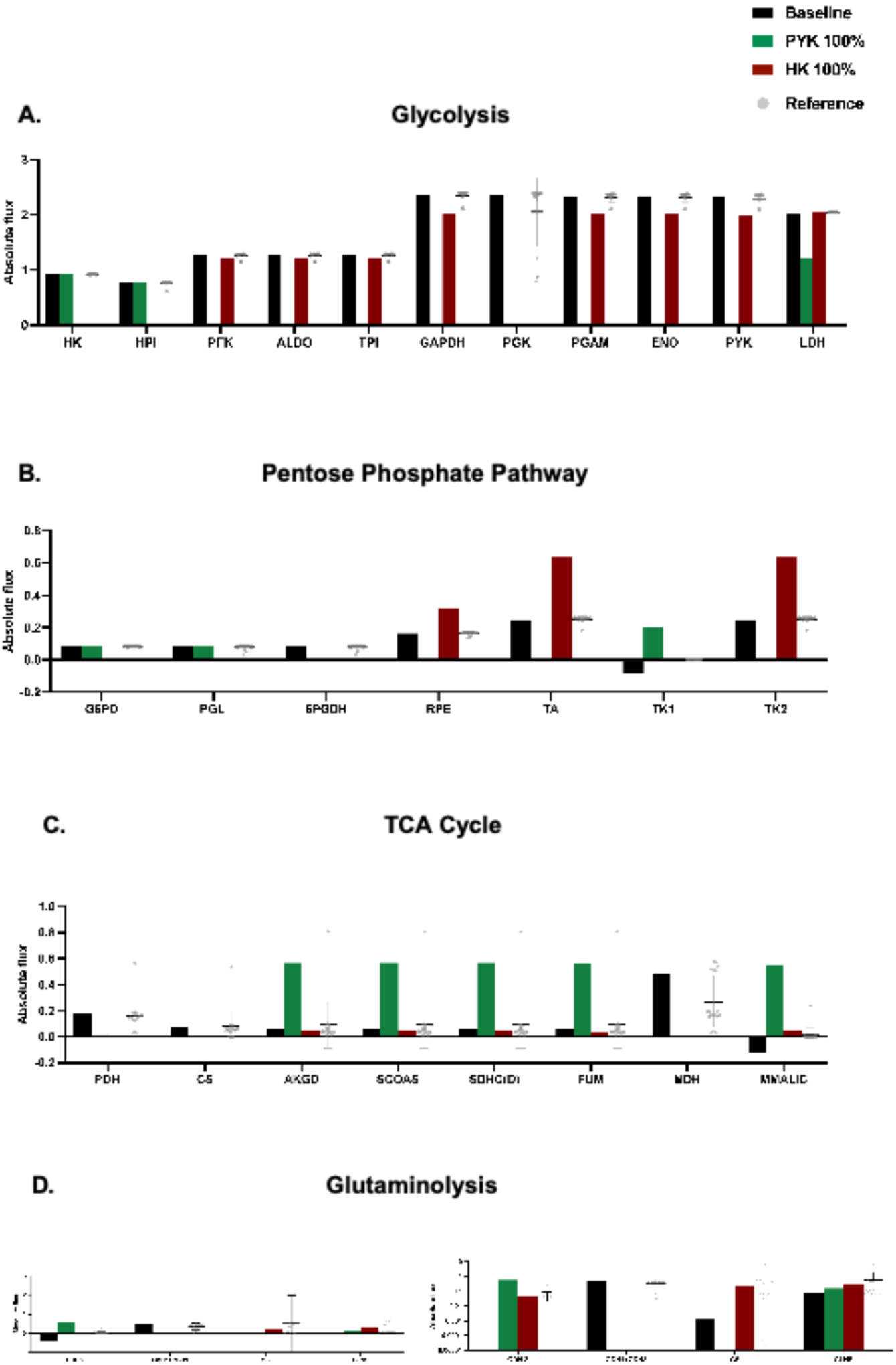
Comparative analysis of the impact of significant perturbations on metabolic pathways for the KRAS_CAF_ condition. Bar plots depicting the fluxes for baseline (no perturbation), PYK 100% knockdown (green), HK 100% knockdown (maroon), and reference (bottom 5th percentile of reactions furthest from the centroid; gray points) for reactions in **(A)** glycolysis, **(B)** PPP, **(C)** TCA cycle, and **(D)** glutaminolysis.

### Pharmacological HK inhibition reduces HK activity in KRAS^MUT^ PDTOs cultured in CAF-CM

Given HK is an enzyme of interest, we investigated whether experimental inhibition would correlate with our *in-silico* results. While the metabolic model includes the general form of the HK enzyme, we focused on inhibiting hexokinase 2 (HK2) using 3-bromopyruvate (3-BP), as it has been shown to be a predictor of patient prognosis.^33–36^ HK2 is used as the isoform experimentally due to specific regulatory and kinetic properties that is applicable for studies on cancer metabolism, while the computational model simplifies the HK reaction to a more generic step in glycolysis.^37^ KRAS^MUT^ PDTOs were treated with 50 and 100μM of 3-BP in CRC media or CAF-CM generated from autologous CAFs. Hexokinase activity was measured after 24 hours of treatment with 3-BP to confirm inhibition (**Figure 5**). Results showed a significant reduction of HK activity to 28.4% in PDTOs cultured in 100μM 3-BP spiked CAF-CM only.

**Figure 5:**
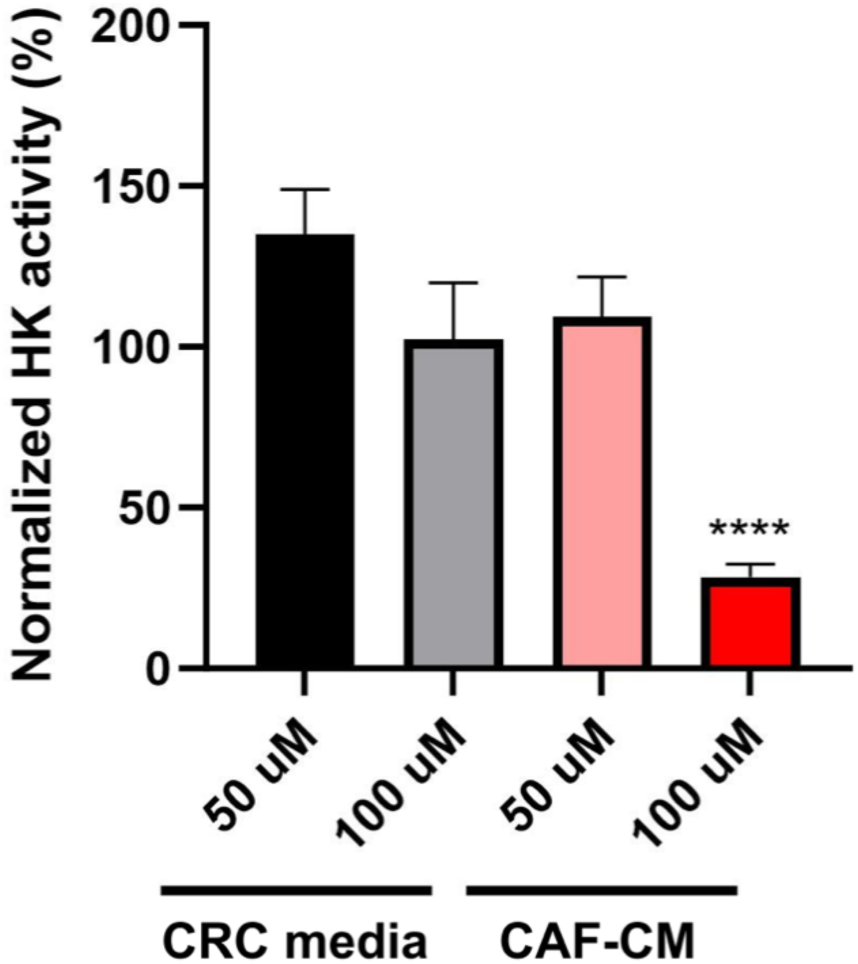
Hexokinase activity in PDTOs treated with 3-BP inhibitor. KRAS^MUT^ PDTOs (000US) treated with 3-BP for 24 hours in CRC media and CAF-CM to measure inhibition of HK activity. Percentage of HK activity is normalized to the respective untreated conditions (CRC or CAF-CM media). \*\*\*\**p*<0.001

### HK inhibition alters metabolic activity and cell viability in PDTOs cultured in CAF-CM

FLIM has been a widely used imaging platform to measure metabolic signatures and drug responses within biological samples;^38–44^ however, to a lesser degree in PDTOs given the recent adoption of this model system in cancer research and the complex 3D structures. The fluorescent lifetime of NAD(P)H was measured within our PDTOs, and phasor analysis was performed by taking the Fourier transform of the fluorescence emission of each pixel of the image.^27,45^ We measured changes in metabolic signatures and quantified the fraction of bound NAD(P)H. An increased fraction bound of NAD(P)H corresponds to a metabolic shift towards oxidative phosphorylation (OXPHOS) and a decrease of fraction bound of NAD(P)H is indicative of a shift towards glycolysis (GLY).^28,39,40,46^

We investigated the metabolic changes in PDTOs during treatment. Initially, PDTOs were a multi-kinase inhibitor, and stained with a vital dye, DRAQ7, to detect dead cells (**Supplemental Figure S10**). Metabolic shifts towards OXPHOS were observed and co-registered with positive staining of DRAQ7, indicating that in these PDTOs, a shift towards OXPHOS correlated with cell death in response to drug treatment. In CRC media conditions treated with various concentrations of 3-BP, there was no significant shift in PDTO metabolic signatures detected by FLIM (**Figure 6A-B**) However, in CAF-CM conditions treated with 3-BP, the metabolic signature shifted significantly towards OXPHOS (increased fraction bound of NAD(P)H, or the ratio of NAD(P)H/NAD(P)+) at the 80 and 100μM drug concentrations, with a concurrent increase in DRAQ7 fluorescence signal detected (indicating regions of cell death). **Supplemental Figure S14** depicts the calculated percent of pixels with positive DRAQ7 compared to the total organoid. We have previously shown a decrease in organoid size correlates with cytotoxicity^47^. As seen in **Figure 6A**, the PDTOs treated with 80 and 100µM 3-BP were observed to be smaller. It is important to note that a decrease in organoid size due to highly effective treatments, such as STA, may skew the %DRAQ7 calculations. As cell death progresses, the DRAQ7 signal could diminish, leading to an underestimation of organoid death within this condition. Therefore, we confirm changes in organoid viability using multiple modalities.

**Figure 6:**
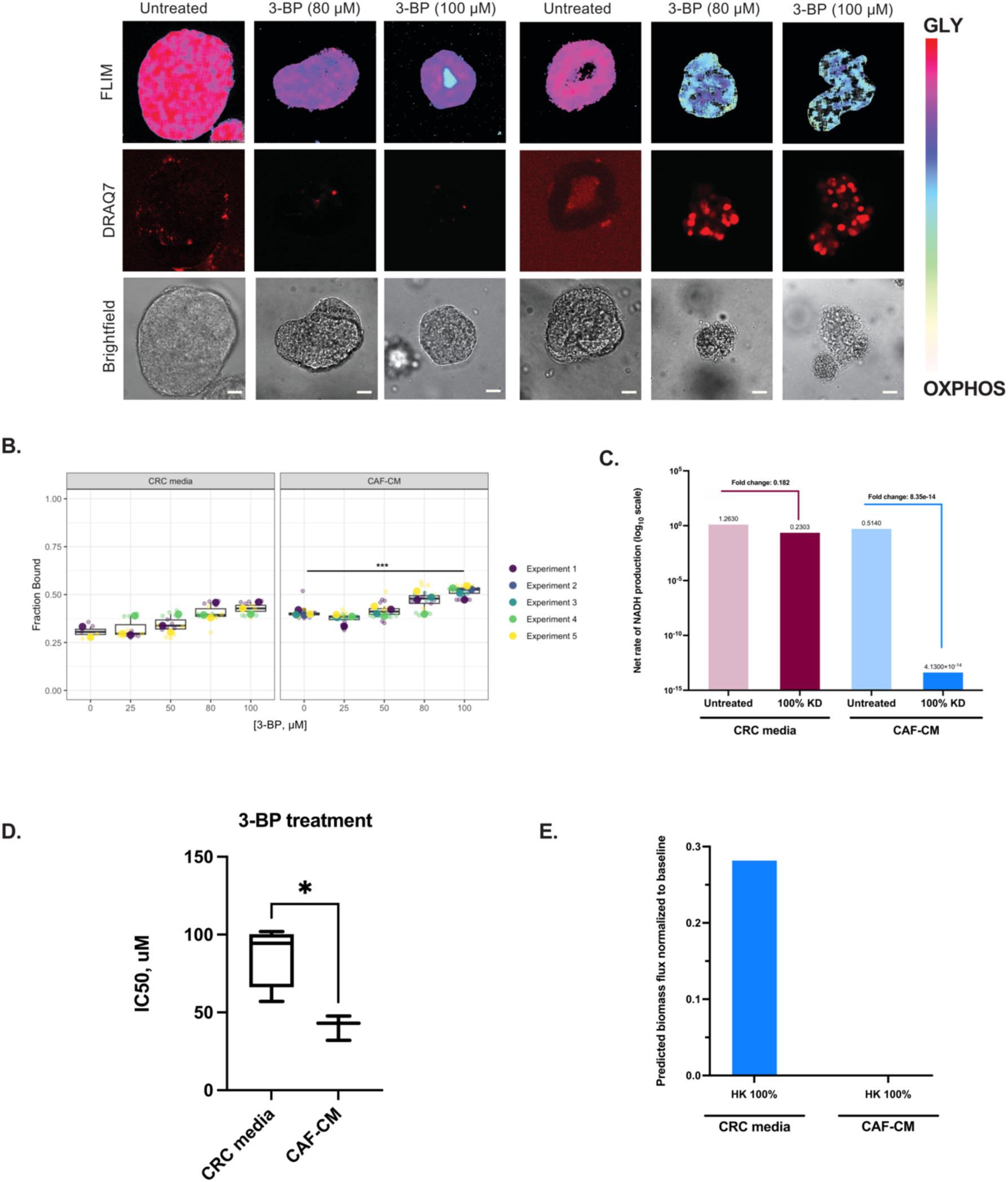
HK inhibition alters metabolic activity and cell viability in CAF-conditioned PDTOs. **(A)** Representative FLIM images and DRAQ7 staining of KRAS^MUT^ PDTOs (000US) in control CRC media or patient-matched CAF-CM after 72 hours of 3-BP treatment. Images are of a single z-slice; scale bar is 20 μm. **(B)** Quantified metabolic signature values comparing PDTOs in CRC media and CAF-CM conditions with 3-BP treatment. For CAF-CM conditions n = 2-3 (2 for 0uM and 3 for 25-100uM). For CRC conditions: n= 4. ***p<0.001**(C)** Model-predicted metabolic signature values comparing cells in CRC media and CAF-CM with HK knockdown on a log_10_ scale. The annotations on the figure indicate the fold-changes, relative to the baseline (untreated) case. **(D)** Calculated IC_50_ values from dose response curves of PDTOs treated with 3-BP in CRC media or CAF-CM. *p<0.05 **(E)** Model-predicted biomass flux for full knockdown of HK in CRC media and CAF-CM normalized to baseline (no model perturbation).

Since FLIM provides a measure of glycolysis and oxidative phosphorylation, we returned to the computational model to calculate and compare the total GLY and OXPHOS fluxes. To reproduce the observed experimental measurements, we defined OXPHOS in the computational model as the net rate of NADH production. Oxidative phosphorylation leads to ATP synthesis using electrons from NADH and FADH2, combined with oxygen. Since our model only considers the cofactor NADH, we calculated the net flux of reactions that produce and consume NADH with and without HK knockout for the CRC media and CAF-CM conditions. This allows for a relevant comparison to the experimental *in vitro* results. We found that the results from the *in silico* analysis align with our experimental findings. Specifically, in both the CRC media and CAF-CM conditions, the model shows that there is a decrease in the net rate of NADH, but with a more substantial reduction in the CAF-CM condition (**Figure 6C**). Namely, the fold change from the untreated condition to the knockdown condition for KRAS_CRC_ is 0.18, while the fold change for KRAS_CAF_ is 4.1×10^-14^.

In addition to the metabolic imaging readouts, we incorporated a cell viability assay, Cell-tier Glo (CTG), based on intracellular ATP levels. The IC_50_ values of PDTOs treated with 3-BP in the CRC or CAF-CM conditions were 83.96 μM and 40.93 μM, respectively (**Figure 6D and Supplementary Table S1**), suggesting CAF-CM renders PTDOs more sensitive to HK inhibition. We again returned to the model to generate calculations that can be directly compared to the experimental measurements. Specifically, we considered the predicted biomass production rates from our model simulations with HK knockdown. The predicted biomass flux can be used to represent cell proliferation, thus decreased biomass flux indicates less proliferation. Zero biomass flux indicates no growth, which could be due to death and birth rates being equal or drugs inducing a cytostatic effect rather than cytotoxic. As expected, the predicted biomass flux was significantly lower for CAF-CM compared to CRC media conditions (**Figure 6E**), showing agreement between experiments and computational modeling predictions.

Additionally, we examined whether this effect was exclusive to HK inhibition or if it could be attributed to a broader stress response by testing clinically relevant chemotherapy drugs, including SN-38, an active metabolite of irinotecan, and 5-fluorouracil (5-FU). SN-38 treatment in the CRC media condition showed a shift towards OXPHOS in a dose-dependent manner after 72 hours of treatment and a positive DRAQ7 staining (**Supplemental Figure S11A-B**); however, 5-FU did not show this metabolic effect. When grown in CAF-CM, fraction bound values also shifted significantly towards OXPHOS at higher concentrations of SN-38, and no change was seen when treated with 5-FU. IC_50_ values from dose-response curves of PDTOs in CRC media conditions treated with SN-38 or 5-FU were calculated to be 0.01 μM and 1.3 μM, respectively (**Supplementary Table S1, Supplemental Figure S11**). In CAF-CM, the average IC_50_ values of PDTOs treated with SN-38 and 5-FU were 0.005 μM and 2.31 μM, respectively. It is important to note that DRCs with 5-FU treatment showed that the IC_50_s for PDTOs in CAF-CM (2.03 μM) were significantly lower compared to when cultured in CRC media (23.93 μM); however, among the six total (three in CAF-CM and three in CRC media) DRC experiments for 5-FU, we never obtained an upper asymptote as defined by Sebaugh’s^48^ bendpoint formula. Diluting 5-FU enough to obtain an upper asymptote was not feasible, so we were only ever able to obtain at most two concentrations with a percent response higher than 50%. Comparison between these IC_50_s with Welch’s *t*-test may be inaccurate as seen in **Supplemental Figure S12C** where their confidence intervals overlap. Overall, treatment with the chemotherapy agents did not result in a differential metabolic shift between CRC and CAF-CM conditions, unlike what was observed with 3-BP treatment.

## Discussion

Metabolic reprogramming of cancer cells has been linked to cancer progression and drug resistance. For example, KRAS^MUT^ colorectal cancer cells have been reported to have increased activity of glycolysis, resulting in increased glucose uptake and lactate. This metabolic reprogramming is correlated with altered cellular growth, and contributes to invasion, metastasis, and drug resistance.^31,32,49^ The TME has been shown to be a key player in supporting these outcomes. Notably, CAFs have a metabolically dependent relationship with tumor cells to promote the survival of each other.^5,6,49–51^ Given the influence of CAFs, it may be possible to exploit the CRC cell-CAF metabolic interactions to inhibit tumor progression.

We have developed a novel, exploratory pipeline to systematically assess, in a high-throughput manner, the comprehensive network effects of enzyme knockdowns in a colorectal cancer-specific model of central carbon metabolism and CAF crosstalk. We combine our *in-silico* predictions with experimental methods to validate our findings and to better understand the effects of metabolic perturbations. Our integrated computational-experimental pipeline identifies enzymes that would be potential therapeutic targets to exploit CRC cell-CAF metabolic crosstalk. A key innovative aspect of this work is the use of representation learning as a form of dimensionality reduction, as it captures the complexities of our input data (fluxes through the entire metabolic network) and allows for meaningful representation of the data in 2D space. We then utilize a distance-based metric to identify influential perturbations that strongly alter the network flux, thus helping us to prioritize experimental testing. Utilization of flux balance analysis on the constraint-based model adapted by Wang *et al.* provides insight into the mechanisms by which CAFs trigger extensive metabolic alterations in CRC.^22^

Initial heatmap analysis of two conditions, KRAS_CAF_ and KRAS_CRC_, show visible metabolic alterations following enzyme knockdowns. For example, glycolytic knockdowns have a more pronounced impact on many metabolic fluxes compared to others. We can also observe the pronounced effects of inhibiting glycolysis on other pathways, as well as on metabolic transport and exchange. This projection of data in 2D space revealed a clear separation among the four conditions. The KRAS_CAF_ cluster distinctly isolates from the other three clusters, suggesting that CAFs uniquely alter the KRAS^MUT^ cells significantly more than the KRAS^WT^ cells and further supports the impact of CAFs on cancer cell metabolism as previously studied by Dias Carvalho *et al*.^52^ Additionally, a clear separation between WT_CAF_ and KRAS_CAF_ highlights genotype-dependent differences, but still a definite impact of CAFs.

We used Euclidean distance measurements within the clusters to identify metabolic perturbations. The resulting distributions of distances from the centroids for all four experimental conditions approximate a lightly tailed normal distribution, enhancing the statistical power of our method. By focusing on perturbations farthest from the mean, we can identify the influential knockdowns. Consistent with previous experimental and computational research, our model suggests that targeting enzymes such as HK, PYK, and LDH -- especially at higher levels, such as 80% or 100% knockdown -- can significantly inhibit metabolism and CRC cell growth. Moreover, overexpressing these enzymes results in cancer cells with enhanced resistance.^53,54^ Furthermore, certain metabolic perturbations, such as PGAM, ENO, and PYK cluster closer together. This indicates that inhibiting these reactions has a similar effect and suggests that they share regulatory characteristics on central carbon metabolism. For example, these enzymes signify the end steps of glycolysis whose reactions are also bottlenecks for being able to carry flux into the TCA cycle. In addition to the critical role of HK inhibition as a rate-limiting step, G6PD inhibition is also identified as a significant approach to perturb the system. This is consistent with the understanding that G6PD serves as a rate-limiting enzyme in the PPP. Therefore, targeting either the initial or terminal step of central carbon metabolism pathways is shown to have profound effects on metabolic flux, altering the system’s overall metabolic profile in diverse ways.-Overall, our framework effectively pinpoints potential strategic interventions to achieve pre-defined, specific metabolic goals.

To experimentally validate our *in-silico* results, we used a label-free imaging-based approach to observe metabolic shifts in response to perturbations within KRAS^MUT^ PDTOs. High-throughput quantitative experimental data is crucial for validating model predictions. However, a major challenge in the field of cancer metabolism is the ability to recreate complex metabolic interactions and patient-specific metabolism *in vitro*.

Bioengineered cancer models, including PDTOs, offer unique opportunities for drug screening. These models leverage characteristics such as patient-level heterogeneity, high-throughput capabilities, and the ability to incorporate multiple cell types and other components of the TME.^43,50,55^ Importantly, drug screening assays using PDTOs have shown a strong positive correlation in their response with that observed in the patient.^41,56,57^

Conventional assays often employed in these drug studies use end-point measurements of viability based on a heterogeneous mixture of PDTOs. Imaging-based applications provide a complementary method for screening, allowing dynamic measurements of drug response. Coupling confocal imaging with vital dyes and brightfield imaging has demonstrated the ability to capture additional information, including changes in growth rates, heterocellular responses, individual organoid morphology, and discerning whether a treatment is cytostatic or cytotoxic.^47,58,59^ Moreover, FLIM enriches our understanding of drug response by examining metabolic signatures within sub-cellular compartments and quantifying the heterogeneity within the sample.^42–44,55,60^ This label-free imaging method can effectively capture metabolic changes over time. Each pixel of the fluorescence emission signal of free NAD(P)H, an endogenous metabolite, is measured and Fourier transformed. This yields a fit-free representation of the imaging data in the form of a phasor plot.^27,45^ The ratio of bound to free NAD(P)H is correlated to regions of relative shifts in glycolysis and oxidative phosphorylation.^28,40,46,61^ An advantage of using FLIM to quantify drug response is the ability to detect drug effects earlier compared to the use of vital dyes. We observed metabolic shifts towards OXPHOS in PDTOs treated with SN-38 as early as 6 hours after treatment, although DRAQ7 staining was not evident until later time points (**Supplemental Figure S13**).

When employing this experimental workflow, we demonstrated that pharmacological inhibition of HK resulted in significant metabolic and viability changes in the CAF-CM condition, which corroborated our *in-silico* HK knockdown simulations. This aligns with research demonstrating that HK is essential for the survival of cancer cells and significantly impacts tumor progression.^62^ To further exhibit that HK2 was a vulnerable enzyme within the CAF-CM setting, we repeated our experiments with two clinically relevant chemotherapy agents. In contrast to our findings with 3-BP treatment, where only the CAF-CM condition exhibited the highest sensitivity to HK inhibition, PDTOs responded similarly to SN-38 in both the CRC media and CAF-CM conditions, showing a shift towards OXPHOS at the highest drug concentrations. Previous studies have shown treatment with irinotecan in KRAS^MUT^ cells resulted in an increase in mitochondria activity, which supports our measured OXPHOS signature. ^63^ Additionally, it has been shown that treatment of CRC cells with irinotecan can result in a decrease in hexokinase activity^64^, which could explain the viability similarities between the 3-BP and SN-38 treatments. Interestingly, 5-FU, a pyrimidine antimetabolite effective in clinical settings, resulted in a decrease in organoid viability in our measurements, yet there was no apparent impact on the main pathways within central carbon metabolism, as indicated by the lack of GLY-OXPHOS metabolic shifts in either the CRC media or CAF-CM conditions from FLIM analysis. Future work exploring dual targeting of glucose and pyrimidine metabolism may enhance the sensitivity of colorectal cancer cells. Overall, our findings suggest that CRC cell-CAF metabolic interactions are more sensitive to targeting HK2, resulting in an enhanced therapeutic response.

### Limitations

We acknowledge some limitations in our study that are important to consider in future research. Primarily, our model is limited to central carbon metabolism. While insightful, this captures a small percentage of cellular metabolism and leaves out pathways and reactions that could play pivotal roles in, or be affected by, the enzyme knockdowns we analyzed. Excluded reactions also lead to compensatory mechanisms in the metabolic network that we cannot consider. Future studies may utilize a genome-scale model (GEM) that encapsulates the entirety of human metabolic reactions for a particular cell type. This would facilitate a more thorough understanding of metabolic reprogramming. Our model is influenced by environmental factors present in the experimental studies, limiting its universal applicability: the conditions under which our experiments were conducted, specifically the glucose, lactate, and glutamine uptake and secretion rates, greatly shape the predicted fluxes. As such, these findings may not be directly transposable across different cell lines or experimental setups. Future work can consider alternate nutrient sources and growth conditions. Moreover, our current study concentrates on the influence of CAF-secreted factors using conditioned media, with follow-up investigations exploring the potential of physical interactions using co-culture models. Additionally, we recognize the limitations of our selected inhibitor, 3-BP, when experimentally validating our computational findings. Although 3-BP is effective at limiting HK2 activity, it, like many pharmacological agents, may interfere with and suppress other biochemical pathways, potentially leading to unintended off-target effects. Future work can utilize more precise targeting of HK2 activity. Lastly, although the primary target of our project was to experimentally validate the inhibition of HK, we recognize the importance of further investigating and experimentally validating other significant targets in the KRAS_CAF_ condition, such as PYK inhibition. Our findings demonstrated differential effects on central carbon metabolism in response to the inhibition of both early (HK) or downstream enzymes (PYK) in glycolysis. Thus, experimental validation of glycolytic perturbations at later stages holds promise for deepening our understanding and refining therapeutic strategies.

## Conclusions

Overall, we demonstrate the utility of our integrated computational-experimental approach for identifying perturbations that exploit metabolic crosstalk between CRC cells and CAFs. Importantly, we show that the model predictions can be validated experimentally using FLIM and conventional cell viability assays. Our established pipeline is versatile and can be integrated with other constraint-based colorectal cancer models, such as GEMs. The approach is also flexible, allowing the inclusion of additional pathways and facilitating double-enzyme knockdown simulations and combinatorial drug treatments. Thus, our work is a foundation for identifying effective strategies for targeting cancer metabolism.

## Materials and Methods

### Constraint-based modeling

We used an existing model of central carbon metabolism, consisting of 74 reactions (including a biomass growth reaction).^22^ Unsteady-state, parsimonious flux balance analysis (upFBA) was utilized to predict the distribution of fluxes for the network. This method was used due to its advantages over other methods such as traditional flux balance analysis (FBA) and unsteady-state FBA (uFBA).^22^ Namely, FBA relies on the assumption of a steady-state system, where metabolite concentrations remain constant and the environment remains non-transient. However, our study focuses on investigating the impacts of different media on CRC cells in a 24-hour period; therefore, traditional FBA is not suitable for our purposes. Moreover, we have informed our model by incorporating constraints on the metabolite fold-changes in the 24-hour period, as measured using metabolomics (previously described in Wang *et al)*.^21^ While uFBA allows for the inclusion of metabolomics data and can investigate dynamical systems, we opted to use upFBA, which predicts the reaction fluxes by minimizing the total flux through the network. The decision to minimize the overall metabolic flux required for biomass production, as opposed to maximizing it as in traditional FBA or uFBA, is based on the optimality principle^65^, which states that cells aim to maintain a specified growth rate while simultaneously minimizing their metabolic expenditures. By performing upFBA, we are able to predict the flux through all 74 reactions given the measured growth rate (which is set as the biomass production rate) and constraints on the metabolite fold-changes over 24 hours.

We applied the model to simulate enzyme knockdowns, with the predicted flux distributions obtained from flux balance analysis serving as the initial reference point for introducing perturbations (“baseline model”). Gene knockdowns were performed using upFBA as well, as a means to predict the maximal biomass growth rate given a perturbation to the system while still minimizing the overall metabolic flux. Thus, the constraint on biomass growth rate used above in upFBA was removed and a double objective function was utilized. For the full enzyme knockouts, the upper and lower bounds for each reaction’s constraints were set to 0 mM/hr. For partial enzyme knockdowns, only the upper or lower bounds were changed relative to the reference flux in the baseline model without the perturbation. We use a 20% knockdown of an enzyme to illustrate our method. If the corresponding enzyme-catalyzed reaction is predicted to proceed in the forward direction in the baseline model, the upper bound would be set to 80% of the predicted flux without any perturbation. Conversely, if the reaction proceeded in the reverse direction, the lower bound would be set to 80% of the predicted flux. Therefore, this method ensures that the enzyme knockdown simulations are context-specific and considered relative to the metabolic behavior observed in the predicted flux distributions from the baseline model. We consider 70 knockdowns out of 74 possible reactions because we do not perturb the reactions responsible for glucose uptake, lactate secretion, glutamine uptake, and the biomass pseudo-reaction, as these are fixed constraints set in the model. All constraint-based modeling was done in MATLAB R2022a, and we utilized the COBRA toolbox v3.0. Scripts relevant to enzyme knockdown can be found on GitHub: https://github.com/FinleyLabUSC/CRC-Cell-uFBA-Modeling.

### Dimensionality reduction

We leveraged representation learning for dimensionality reduction, facilitating the analysis and comparison of a large set of vectors. Representation learning employs neural networks to transform a wide range of inputs into low-dimensional features. In this work, we specifically used Siamese neural networks (SNN), which are trained on all inputs using the same network architecture.^30^ As a result, the projected low-dimensional vectors can be directly compared to each other as a measure of similarity, which is our goal. Furthermore, this approach enables comparison of the input data without the need for additional data processing, such as normalization.

We consider a vector of reaction fluxes, each 1x74 in size, for individual enzyme knockdowns. These vectors represent the predicted flux through 74 metabolic reactions and signify the systemic impact of these knockdowns. Our simulation includes five levels of partial enzyme knockdowns (20%, 40%, 60%, 80%, and 100%) across four different conditions (WT_CRC_, WT_CAF_, KRAS_CRC_, and KRAS_CAF_). Consequently, we have generated a large dataset comprising 20 sets of 70x74 vectors from this metabolic modeling (1400 total perturbations). We then employ a neural network that interprets these input matrices as if they were images, with each matrix value being analogous to a pixel’s intensity in an image. The network is designed to discern subtle patterns within these ‘images,’ especially focusing on features around the diagonal, which correspond to the enzyme knockdowns. The neural network’s capability to discern subtle patterns is influenced by how the data is structured and presented. Large matrices containing all 70 knockdowns for a given condition and knockdown level are fed into the neural network, allowing it to learn from the interactions between the rows and columns of the specific input file. This is critical for creating a meaningful mapping in 2D space because the network trains on these points in the context of the larger matrices, capturing relationships within the rows and columns and optimizing the embedding space based on a complex set of features in relation to one another. This can be compared to the method by which a sophisticated car recognition system processes images of vehicles. Just as this system analyzes and learns from various visual aspects of a car (such as shape, color, etc.) to differentiate different types of cars, the neural network examines the matrices representing enzyme knockdowns as if they were intricate patterns in an image. Each ‘pixel’ or data point in the matrix is weighed and considered in the context of the full ‘image’, or the complete matrix of knockdowns, to distinguish the overarching structural features that denote similarity or dissimilarity, in the way that cars are identified in this way based on various characteristics. Through this method, the network will be able to construct a low-dimensional embedding that captures the structural features of the data, learning a representation that intrinsically captures similarities in the inputs. By computing the distance between these inputs in 2D space, we can determine the similarity between the inputs, yielding a single value that reflects the complex interrelations among the model’s output features.^30^ As such, the dataset is better suited for representation learning, compared to other methods.^23^

Other methods of dimensionality reduction do not produce new, learned representations of the data, but rather project the original features into low-dimensional space comprised of the original input vectors, such as in t-SNE. However, neural networks have been found in numerous previous works to reduce the dimensionality of images more effectively compared to methods such as t-SNE.^66^ This is in large part due to complexities within large sets of vector data that cannot be measured using predefined features. Additionally, other methods of dimensionality reduction, such as PCA, are also limited in their ability to faithfully represent the data. For example, PCA fails to capture complexities in large, non-linear data sets since it only finds linear combinations of the original features.^67^

In our approach, the SNNs are trained on the full input set through conventional triplet loss training. Note that we do not consider a test set, as we are not using the trained SNN for prediction purposes. We employ this supervised form of learning because unlike unsupervised learning, it can capture local and global features of high-dimensional datasets.^68^

The neural network begins with layers of neurons, each with an associated weight initially set to random values from a uniform distribution. As data moves through these layers, it is transformed, reducing its dimensionality and capturing key features. The network learns to represent the data by adjusting these weights, guided by bias vectors, through each training iteration. This learning is driven by a loss function, which evaluates how accurately the network clusters similar data points together while separating distinct ones through the calculation of their Euclidean distances.^23^ By optimizing this function, the network iteratively improves its positioning of points in a lower-dimensional space, ensuring that similar data points are closer together. Backpropagation is central to this process, adjusting the bias and weight values so that, with each iteration, the network better captures the underlying data structure.^30^ As training progresses, the network refines its ability to represent relationships between points in a 2D embedding, with the distances between them reflecting both similarities and differences within the same condition or across different conditions. This interpretability in point distances is often more meaningful than in other dimensionality reduction techniques, like t-SNE or UMAP, which may not preserve these relationships as effectively. As we input each vector of fluxes into the neural network, it generates a corresponding triplet, which is then processed further. This triplet consists of an anchor (the original data point), a positive data point (a data point in the set of inputs that is similar to the anchor), and a negative data point (a data point in the set of inputs that is different from the anchor). Since we are interested in understanding differences between the projected data points, we calculate the Euclidean distance between points, traditionally used in triplet loss. This conveys the distance between projected data as a measure of dissimilarity or similarity (**Figure 7A-C)**. It is important to note that the locations of projected points are influenced by the network and type of training, and therefore, the actual distances do not correspond to a physical quantity. Please see Supplemental Material for justification of use of Euclidean distance in the 2D space. Scripts relevant to representation learning can be found on GitHub: https://github.com/FinleyLabUSC/CRC-Cell-uFBA-Modeling.

**Figure 7:**
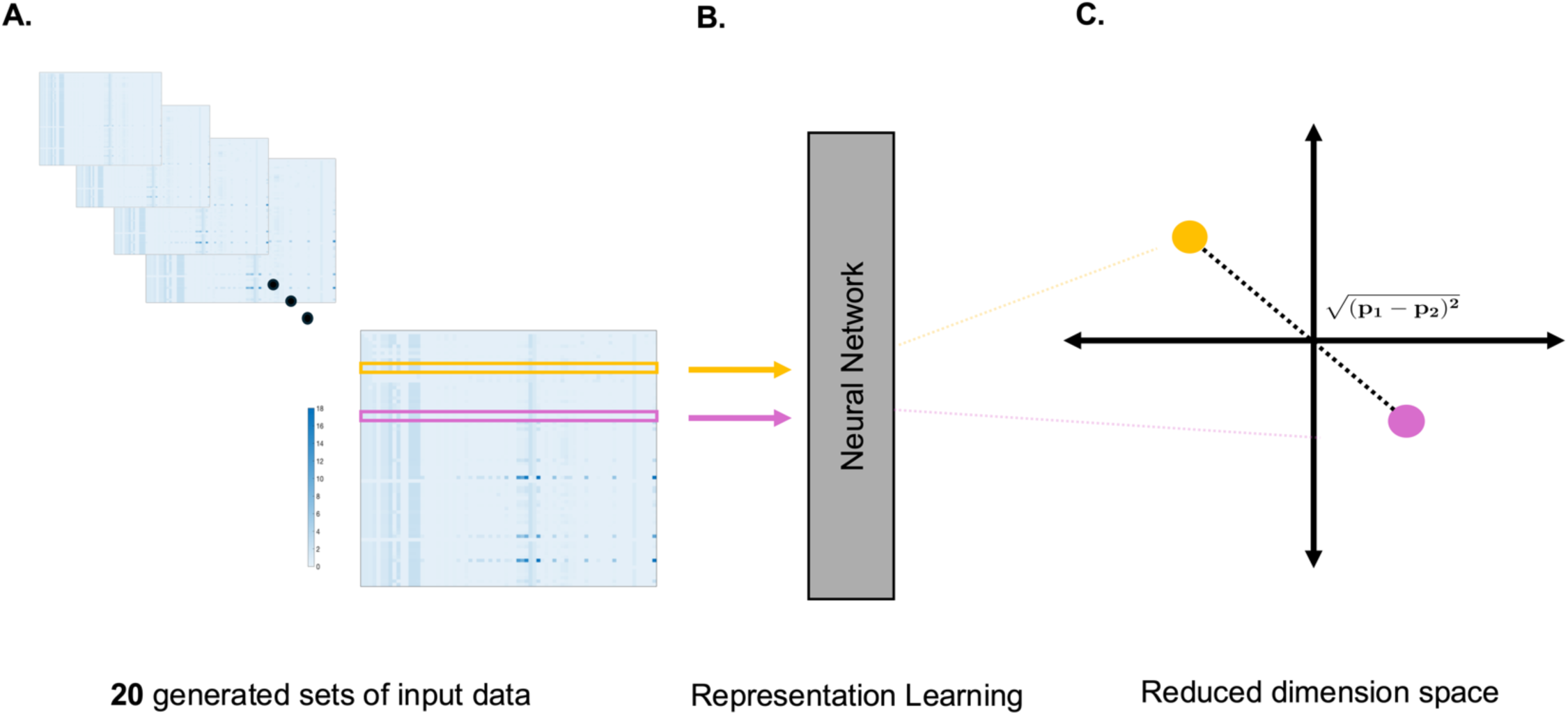
Dimensionality Reduction Pipeline. **(A)** Our simulations resulted in 20 input matrices representing 5 levels of partial to full enzyme inhibition across 4 distinct cell culture conditions. **(B)**: Each row of these matrices was input to a neural network, which iteratively learned the similarity and dissimilarity among data points, projecting them into a 2D space. **(C)**: Through multiple simulation iterations, the algorithm refined point placement to enhance clustering of similar data points, with the distance between points indicating the level of similarity.

### Cell culture conditions

#### Maintenance and cell line verification

PDTOs and CAFs were generated from CRC tumor resections received from USC Norris Comprehensive Cancer Center Translational Pathology Core according to Institutional Review Board (Protocol HS-06-00678) approval as outlined previously^47,59,69^. All patients gave their informed consent before they were included in the study. Organoids were sequenced by the USC Molecular Genomics Core using Illumina’s TruSight Oncology 500 next-generation sequencing platform to evaluate cancer biomarkers present (**Supplementary Table S3**). Immunostaining of PDTOs was also done to confirm cellular composition of organoids (**Supplementary Figure S15**). Characterization of the CAFs was done by our group and previously published^70^.

PDTOs and CAFs were grown in 5% CO_2_ and 37°C and maintained in CRC media, consisting of Advanced DMEM/F-12 media (Thermofisher Scientific, Waltham, MA; 12634010) and the following supplements (at their noted concentrations and solvent) outlined in Table 1:

**Table 1:**
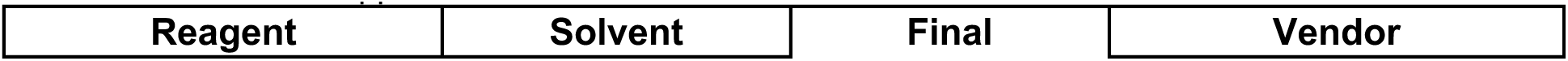
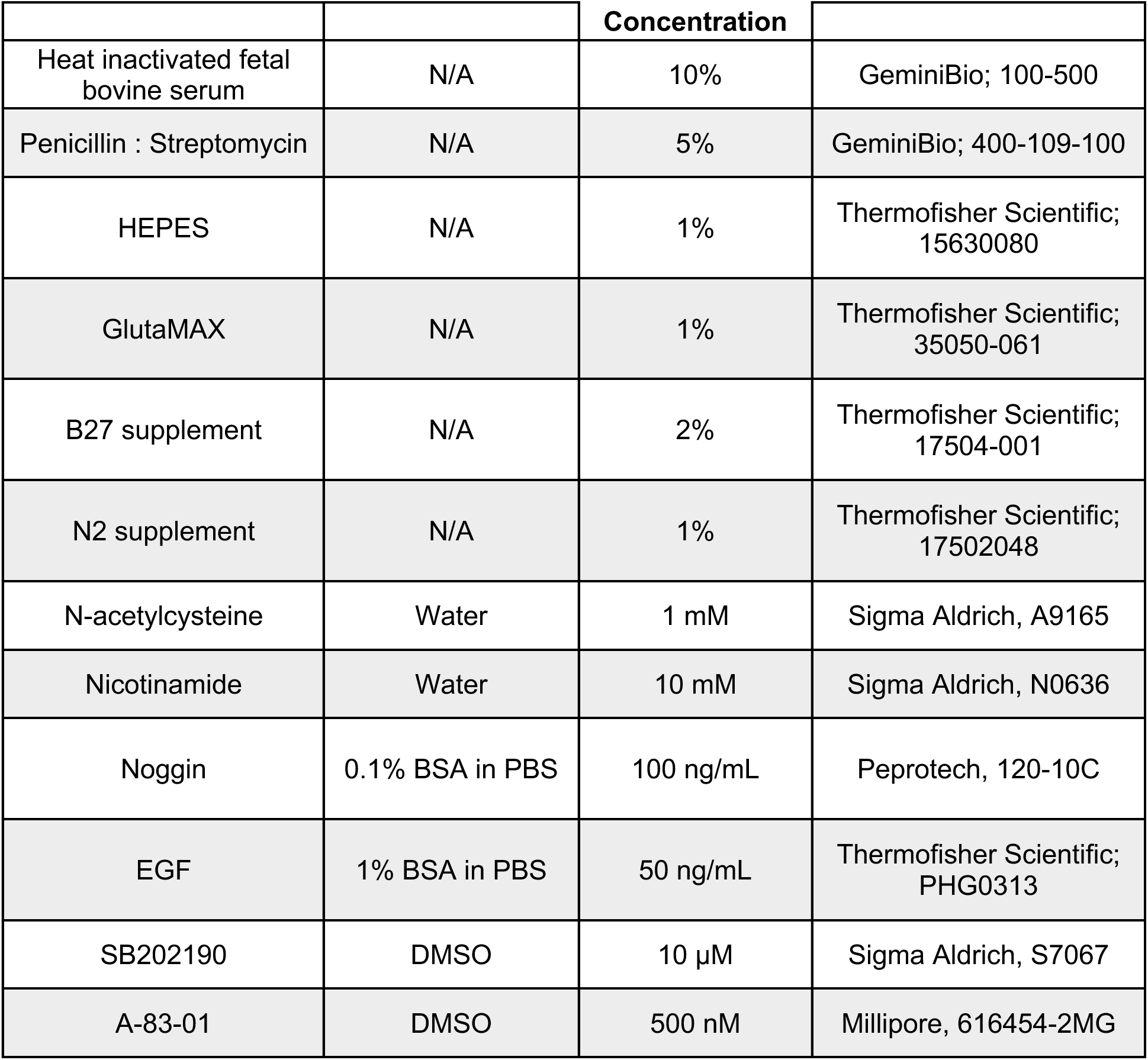
CRC media supplements.

PDTOs were maintained by embedding in Cultrex Reduced Growth Factor Basement Membrane Extract, Type 2 (BME; R&D Systems, Minneapolis, MN; 3533-001-02) and droplets of 60μL were plated onto 24-well plates. The plates were allowed to slightly gel at room temperature for 1 minute before flipping over and placed into the incubator for 15 additional minutes for full gelling. CRC media was then gently added to the wells to maintain the cultures.

#### Organoid immunostaining

The organoid samples in BME were fixed with 4% paraformaldehyde at 4^0^C for 20 minutes, followed by three DPBS rinses. Fixed samples were then permeabilized with 0.1% Triton X-100 for 20 minutes at room temperature. Blocking was done with 2% BSA solution to prevent the non-specific binding for 20 minutes. Cultured organoid samples were incubated with primary antibody cocktail at 4^0^C overnight. The secondary antibody cocktail was used to stain the samples the next day for two hours. The formulas of the antibody cocktails are listed in **Table 2**. The samples were rinsed with DPBS three times between each step.

**Table 2:** Formula of antibody cocktails for immunostaining.

| Primary antibody cocktail |  |  |
| --- | --- | --- |
| Antibody | Dilution | Vendor |
| anti-CK20 | 1:200 | Abcam; ab854 |
| anti-vimentin | 1:200 | Abcam; ab92547 |
| Secondary antibody cocktail |  |  |
| Antibody | Dilution | Company |
| goat anti-mouse | 1:200 | ThermoFisher; A-21424 |
| goat anti-rabbit | 1:200 | ThermoFisher; A-11008 |
| DAPI | 50 ng/mL | Sigma-Aldrich; D9542-5MG |

#### Assay set-up

PDTOs were dissociated into single cells when prepared for imaging and CellTiter-Glo (CTG; Promega, Madison, WI, G9681) cell viability assays. 500 μL Gentle Cell Dissociation Reagent (STEMCELL Technology, Cambridge, MA; 07174) was added to each well, followed by scraping of the organoid/BME droplets with a P1000 pipette tip. This cell suspension was then rocked at 4°C for 30-35 mins in a 15 ml conical tube. Organoid suspensions were centrifuged at 300x*g* for 5 minutes and the BME/dissociation reagent was carefully removed. A 1:1 fragmentation solution of 1X DPBS and TrypLE (Thermofisher Scientific, Waltham, MA; 12605028) plus 1:1000 ROCK inhibitor, Y-27632 (STEMCELL Technology, Cambridge, MA; 72302) was used to resuspend the cell pellet and allowed to incubate at 37°C for 1-2 minutes with physical agitation every 30 seconds. Periodic checks of the cell suspension were done under a microscope to ensure that a majority of the organoids were fragmented. After confirmation that most organoids were dissociated to single cells, the fragmentation solution was quenched by adding 2X volume of CRC media to the cell suspension. The cells were then centrifuged at 300x*g* for 5 minutes and resuspended in CRC media. The single-cell suspension was passed through a 40 μm cell strainer to remove larger organoid fragments. Live cell counts were measured using the TC20 automatic cell counter (Bio-Rad, Hercules, CA). The required amount of cells/assay were suspended in BME with 10% collagen I (Sigma Aldrich, St. Louis, MO; C3867, stock concentration 2.5 mg/mL) to a concentration of 100,000 cells/mL. 10 μL of the cell suspension was plated as a droplet to each well of a white-wall clear-bottom 96-well plate (Corning, Glendale, AZ; 3903) for CTG assays or a clear-walled glass-bottom plate (Mattek, Ashland, MA; P96G-1.5-5-F) for FLIM imaging. Plates were incubated for 10 minutes at 37°C inverted. 200 μL of CRC media containing 1:1000 ROCK inhibitor Y-27632 (STEMCELL Technology, Cambridge, MA; 72302) was then added to each well to maintain the organoids until drug treatments were initiated.

#### Conditioned media

CAF conditioned media (CAF-CM) was generated when patient derived CAFs were grown in 10 cm^2^ tissue culture dishes to about 70-80% confluency in CRC media. Media was refreshed with new CRC media and allowed to be conditioned by the CAFs for 72 hours before collection and filtering through a 40 μm pore size filter. Collected media was stored at −30°C until assay use.

### Drug treatments and cell viability assays

Various concentrations of staurosporine (STA; Sigma-Aldrich, St. Louis, MO; 569396), 5-fluorouracil (5-FU; Selleck Chemicals, S1209), 7-ethyl-10-hydroxycamtothecin (SN-38; Sigma-Aldrich, St. Louis, MO; H0165), or 3-bromopyruvate (3-BP; Sigma-Aldrich, St. Louis, MO; 16490) were added to their respective wells. For plates reserved for FLIM imaging, DRAQ-7 (Abcam, Cambridge, MA; ab109202) was added to each well at a final concentration of 3 μM. Control samples to demonstrate metabolic inhibition were treated with 100 mM sodium dichloroacetate (Sigma-Aldrich, St. Louis, MO; 347795) and 50 mM 2-deoxyglucose (Sigma-Aldrich, St. Louis, MO; D6134) for 24 hours prior to FLIM imaging.

For cell viability assays, PDTOs were cultured for 4 days in CRC media before the addition of any drugs, as described above. After 5 days of treatment, CTG assays were conducted as described by the manufacturer. The CTG assay is used to calculate the percentage of cells within the population which remain viable at a range of drug doses. The drug concentration at 50% viability is termed “IC_50_” and is used as a reference to compare drug sensitivity across different treatments. A lower IC_50_ indicates that a lower drug concentration is required to induce cell death in that sample. For FLIM experiments, organoids were grown for 7 days and then treated with drugs for 6 or 72 hours prior to imaging.

### Fluorescence lifetime imaging microscopy (FLIM) and data analysis

FLIM imaging was done at the Translational Imaging Center (TIC) at the University of Southern California (USC) and at the Ellison Institute of Technology (EIT). For images taken at the TIC, a Zeiss 780 inverted confocal microscope (Carl Zeiss, Jena, Germany) was used. 256x256 pixel sized images that were taken with a Plan-Apochromat M27 20X/0.8 NA (Carl Zeiss, Jena, Germany) objective at 12.6 μs pixel dwell time. The PDTO samples were excited with a Ti:Sapphire 2-photon laser (Coherent Chameleon Ultra II) set at 740 nm for NAD(P)H excitation. The emission was split into two channels with a 590 nm long-pass filter and each channel was further filtered at 460/80 nm for NAD(P)H. External photomultiplier tube detectors (Hamamatsu, Japan R10467U-40) were used to detect the emission for each channel. An A320 FastFLIM FLIMbox (ISS Inc., Champaign, IL.) was used to collect the frequency domain lifetime of NAD(P)H with VistaVision software (ISS Inc., Champaign, IL). An averaging of 4 frames per z-slice was done to collect enough photons for FLIM analysis. For confocal images on the Zeiss 780, 512x512 images were taken with a 633 nm excitation for DRAQ7.

FLIM imaging performed at EIT was done on an Olympus FV3000 microscope with a UPLXAPO 20X/0.8NA objective, which was upgraded with an A320 FastFLIM FLIMbox (ISS Inc., Champaign, IL.) add-on and hybrid PMT external detectors (Hamamatsu, Shizuoka, Japan). Samples were excited at 740 nm with a Ti:Sapphire 2-photon laser (Spectra-Physics, Mountain View, CA) at 15-20 mW and the emission was passed through a 458/64 dichroic for NAD(P)H collection. VistaVision software (ISS Inc., Champaign, IL) was used to collect the data. 256x256 images were taken with a pixel dwell time of 10 μs/pixel. An averaging of 10 frames per z-slice was done to collect enough photons for FLIM analysis. 512x512 confocal images for DRAQ7 vital dye staining were taken with 640 nm excitation.

Both systems were calibrated in a similar fashion with a solution of coumarin-6 in 200 proof ethanol (known lifetime of 2.5 ns). Collection of the FLIM data was done until the average photon count was about 100 counts/pixel. Organoids of 100-120 μm were manually selected for imaging to ensure that oxygen diffusion limits were not a contributing factor. Three z-slices of each organoid were taken, where the middle slice of the z-stack was the center of the organoid. Image files were exported as .R64 files through VistaVision and analyzed with SimFCS software (Laboratory for Fluorescence Dynamics, University of California, Irvine). It is important to note that the figure images displayed are a single z-slice. Thus, the FLIM signal is only efficiently collected at the in-focus planes although background out-of-focus regions can be seen in our brightfield images.

### Hexokinase (HK) activity assay

Dissociated PDTO single cells were mixed with BME and seeded in a 24-well plate at 200,000 cells/mL. 60 uL of the BME cell suspension were seeded in each well and incubated inverted at 37°C for 10 mins to allow for BME gelling. Organoids were grown for 7 days in CRC media before being treated with CRC media or CAF-CM spiked with 3-BP. After treatment for 24 hours, organoids were digested with a gentle dissociation buffer and rocked for 40 minutes at 4°C. Cells were collected by centrifugation at 300x*g* for 5 minutes. BCA protein assay (Thermofisher Scientific, Waltham, MA; 23225) was conducted to normalize protein concentrations between all samples and hexokinase activity was measured according to the manufacturer’s protocol (Abcam, Cambridge, MA; ab136957), using 10 μg of protein per condition. Readings were collected on a Biotek Synergy Neo2 plate reader (Agilent, Santa Clara, CA) every 5 minutes for 3 hours to capture the window of optimal reaction readings for initial and final time points.

### Statistical analysis

Statistical analysis was performed using R in the RStudio environment.^71^

For FLIM imaging, fraction bound was calculated for three Z-slices of each organoid, then averaged together to produce one summary fraction bound value for each organoid. To account for the variation between organoids within each well we used a mixed effect model constructed with the lmer() function from version 1.1-35.1 of the lme R^38^ package. We constructed mixed effect models for each test drug with an interaction term for drug dose and media type to examine how the effects of each drug changed when adding CAFs.

To examine the action of the drugs at individual doses, we corrected for differences between experiments with batchma’s adjust() function,^72^ checked normality using the check_normality function from the performance package, and conducted an ANOVA test using stats’s aov() to compare the fraction bound between doses of 3-BP for CAF and CRC media types.

Data from cell viability CTG experiments were used to construct dose response curves (DRCs) for all experimental conditions. Drug conditions were normalized to within-plate negative control (untreated) wells and positive control wells (STA treated). These controls were used to calculate percent relative viability using the following formula:

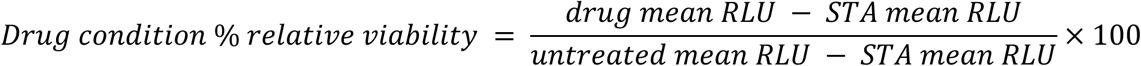

Where drug mean RLU corresponds to the mean relative luminescence units (RLU) over six technical replicate wells in a single plate. STA mean RLU corresponds to the mean RLU of the six technical replicate wells of STA; and untreated mean RLU corresponds to the mean RLU of the six technical replicated wells of the untreated condition. Individual wells were submitted to image QC and excluded if microscopy images showed evidence of debris or poor plating.

The drug condition % relative viability results from a single plate were examined together and used to model a DRC using R Studio^73^. The *logistic()* function from version 3.0.1 of the *drc*^74^ R^71^ package was used to determine which log-logistic regression model fit each curve. Most compounds produced a four-parameter curve, which were subsequently produced (after model selection) using the LL.4() function from the same version of drc. SN-38, however, produced a five-parameter asymmetric curve in all biological replicates. Dose response curves for SN-38 were subsequently formulated using *logistic()*.

After modeling the DRCs for each biological replicate, we evaluated them using several quality control (QC) criteria **(Supplementary Table S4**). We calculated Z’Factor^75^ for each biological replicate and used only experiments where this statistic was greater than 0 to ensure that a sufficient window between positive and negative controls existed to accurately examine differences in RLU between drug conditions. We calculated plate

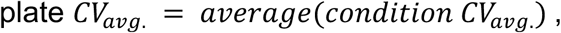

Where

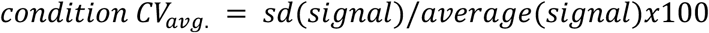

for each compound and concentration.

Iverson *et. al.*^76^ recommends a bound of 20% for homogeneous cell line assays, but PDTOs displayed a more variable signal by condition, so we used a bound of 30%. Absolute IC_50_s were used to compare potency between conditions, so biological replicates with extrapolated IC_50_s were considered to fail quality control criteria. Residual standard error (RSE) was calculated by the *drm*() function from version 3.0.1 of the drc R package. We used a bound of RSE < 20 to eliminate dose response models that may have been made less accurate by residuals. Finally, models with an upper asymptote 95% confidence interval below 55% relative viability were eliminated to ensure that we only examined models with an upper asymptote that is closer to 100% relative viability.

While only biological replicates that passed the above quality control criteria were considered for official analysis, we ensured that the results were the same when all biological replicates were considered. See **Supplementary Table S4** for summary of QC metrics and number of runs for each condition which passed QC. Absolute IC_50_ means were significantly different between replicates cultured with and without CAF-CM when considering all replicates using Welch’s *t*-test (*p*-value = 0.020). When using all replicates, excluding those that did not pass QC criteria, absolute IC_50_s were found to be significantly different using the same test (*p*-value = 0.007).

Scripts for statistical analysis are available on GitHub at: https://github.com/FinleyLabUSC/CRC-Cell-uFBA-Modeling.

## Author contributions

N.T. conducted all metabolic modeling and computational analyses. E.J.F. conducted or directed all experimental sample preparation, data collection, and analysis for FLIM experiments, cell viability measurements, and hexokinase activity assays. FLIM imaging and data analysis was also done by Y.K.H. M.B. collected CTG and FLIM organoid data. Image-based quality control and experimental planning was done by M.D. and S.K. A.C. performed quality control analysis of all CellTiter-Glo and FLIM experiments and calculations of IC_50_ values. HJL provided CRC patient samples and clinical expertise. E.J.F., N.T., S.M.M., P.M., N.A.G., and S.D.F. contributed to manuscript writing and scientific discussions of results. S.M.M. and S.D.F. were responsible for the scientific direction of this work. All authors reviewed and provided feedback on the manuscript.

## Acknowledgements

We would like to thank the members of the Ellison Institute of Technology and Finley Lab at USC for the deep scientific discussions. Special thanks to Scott Valena and Pratiksha Kshetri for cell culture assistance, Nolan Ung for statistical guidance, the Translational Imaging Center at USC for providing us usage of their microscope systems. We acknowledge Yuyuan Zhao for providing immunostained images of the organoids, the USC Norris Comprehensive Cancer Center Translational Pathology Core and the Molecular Genomics Core for the collection of patient tissue samples and targeted sequencing of our patient organoid lines (Norris Comprehensive Cancer Center CCSG grant, P30CA014089). We acknowledge funding support from the USC Center for Computational Modeling of Cancer and the National Cancer Institute Cancer Systems Biology Consortium (U01CA232137).

## Data availability

The datasets and computer code produced in this study are available in GitHub (https://github.com/FinleyLabUSC/CRC-Cell-uFBA-Modeling).

