## Supplemental Material for "Merging Metabolic Modeling and Imaging for Screening Therapeutic Targets in Colorectal Cancer"


### Supplementary Figures

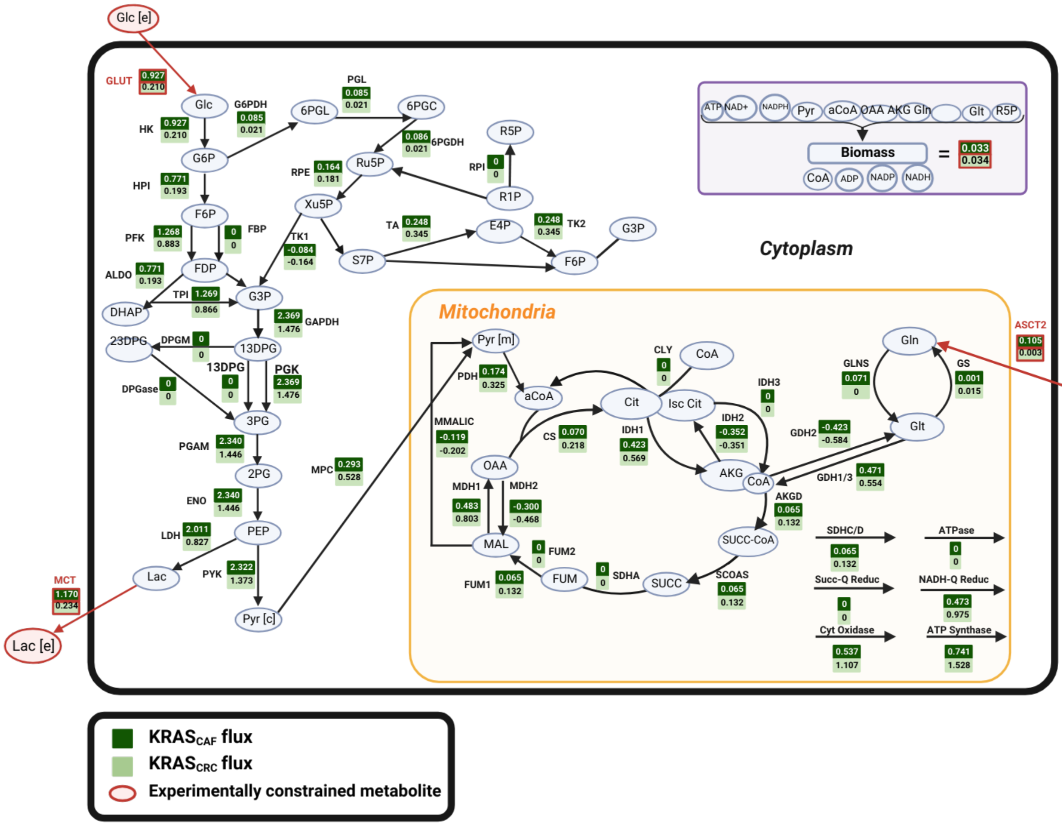


**Supplementary Figure S1: Network of central carbon metabolism depicting the baseline fluxes for KRAS$_{CAF}$ and KRAS$_{CRC}$.** Flux values in red represent experimentally constrained values predefined in the model.

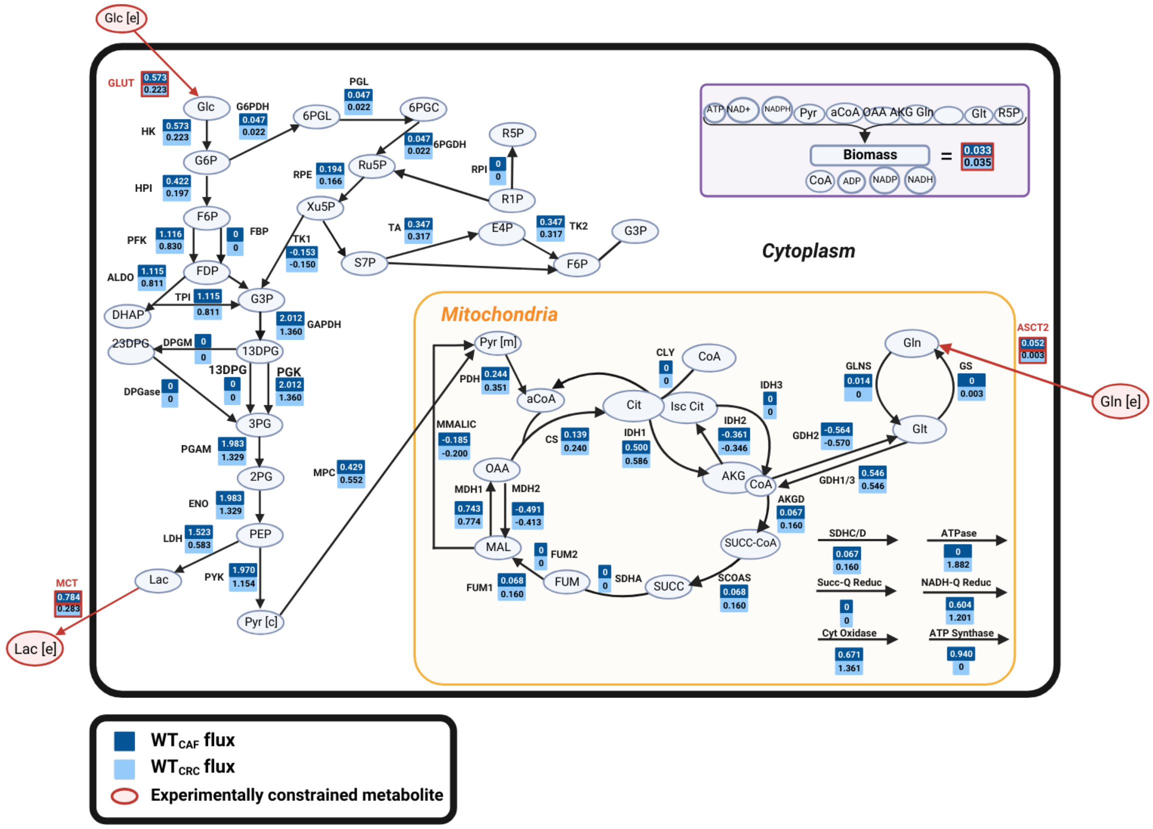


**Supplementary Figure S2: Network of central carbon metabolism depicting the baseline fluxes for $WT_{CAF}$ and $WT_{CRC}$.** Flux values in red represent experimentally constrained values predefined in the model.

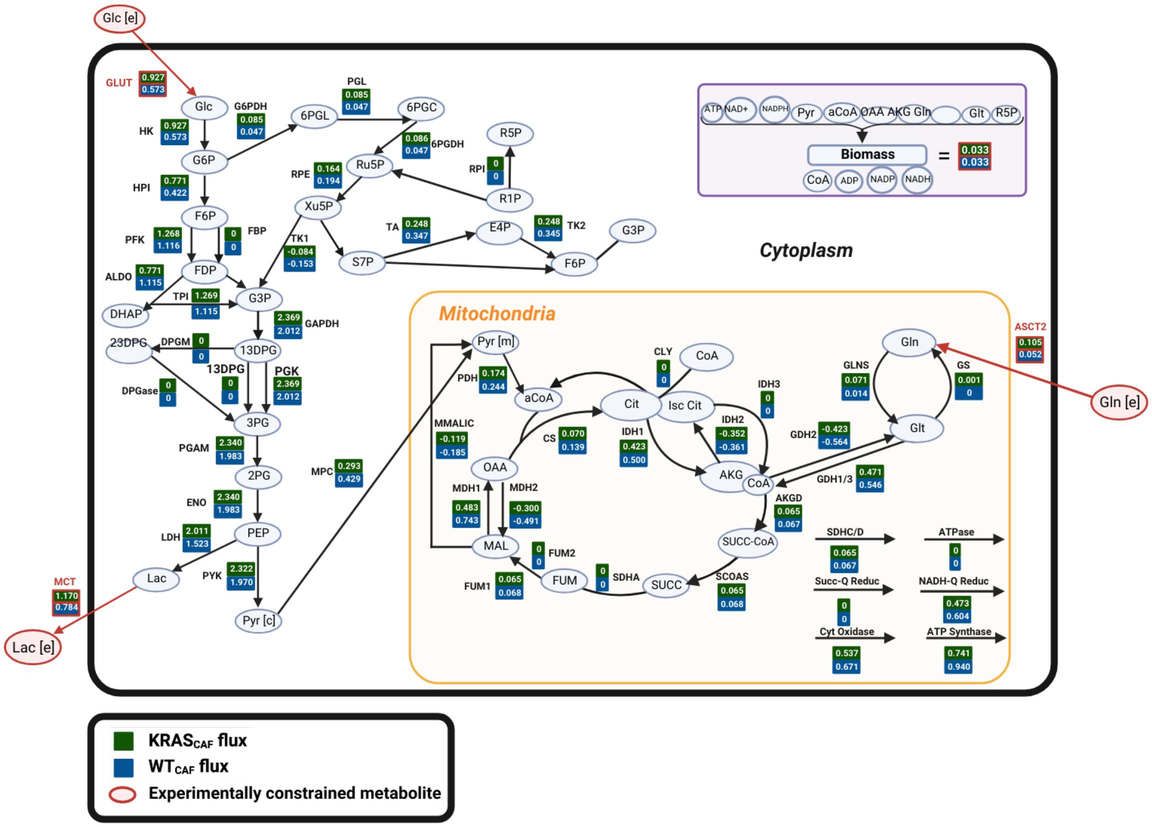


**Supplementary Figure S3: Network of central carbon metabolism depicting the baseline fluxes for $KRAS_{CAF}$ and $WT_{CAF}$.** Flux values in red represent experimentally constrained values predefined in the model.

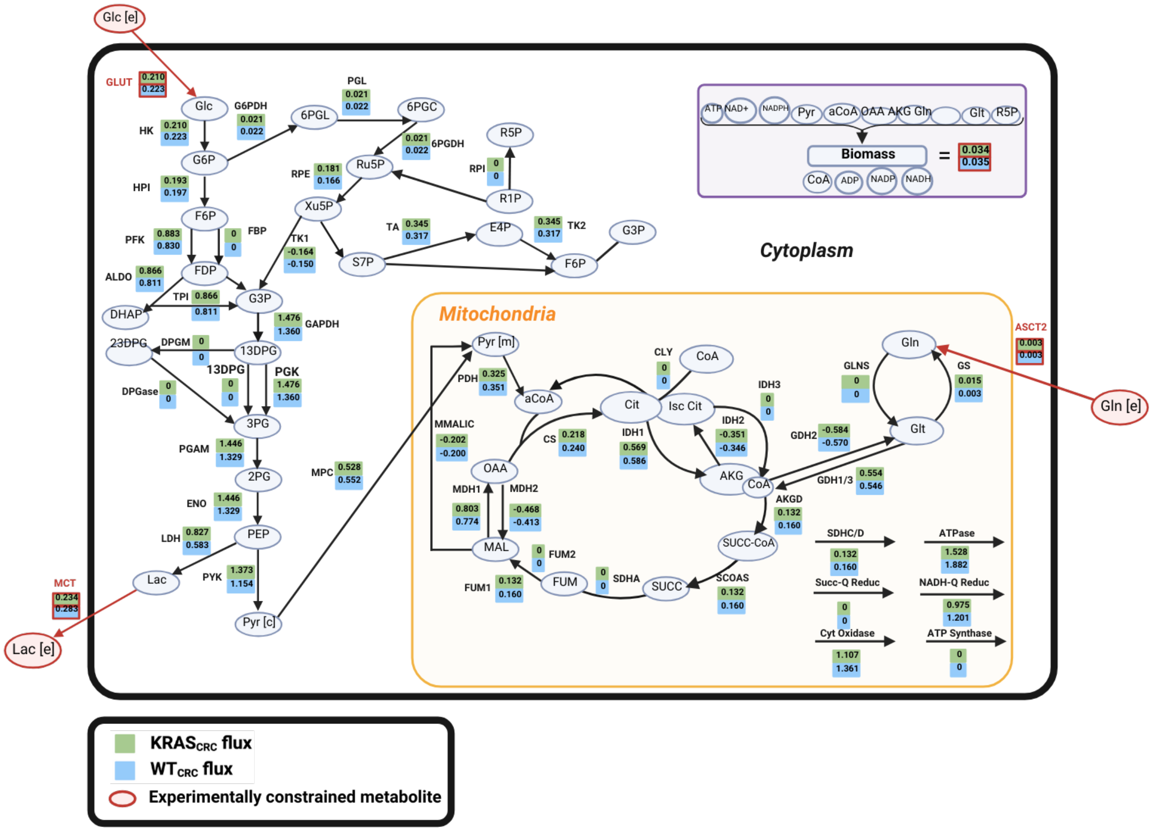


**Supplementary Figure S4: Network of central carbon metabolism depicting the baseline fluxes for $KRAS_{CRC}$ and $WT_{CRC}$.** Flux values in red represent experimentally constrained values predefined in the model.

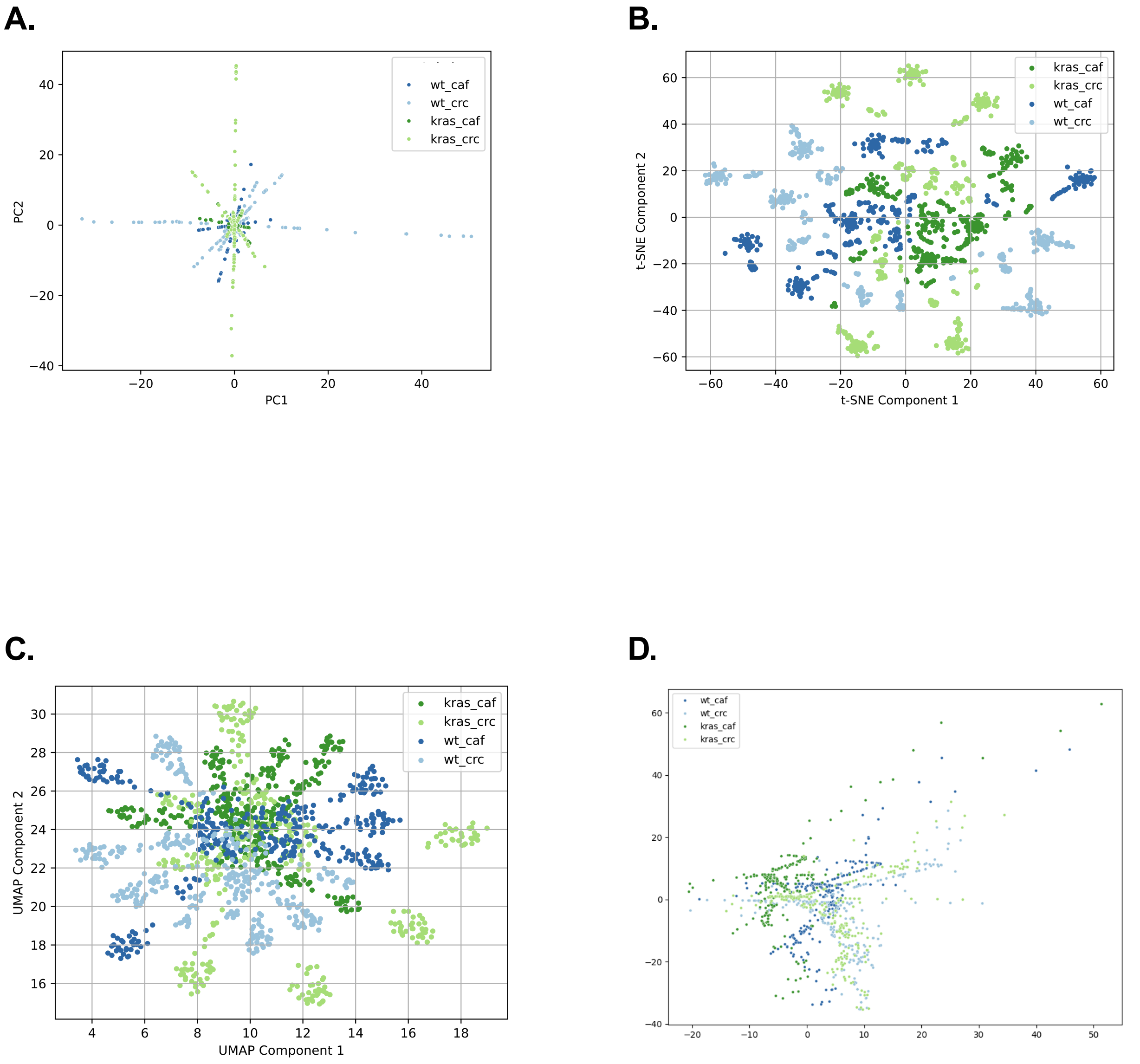


**Supplementary Figure S5: Alternatively used methods of dimensionality reduction. (A)** Principal components analysis (PCA). **(B)** t-distributed stochastic neighbor embedding (t-SNE). **(C)** Unifold maniform approximation and projection. **(D)** Representation learning using unsupervised learning (trained using simCLR).

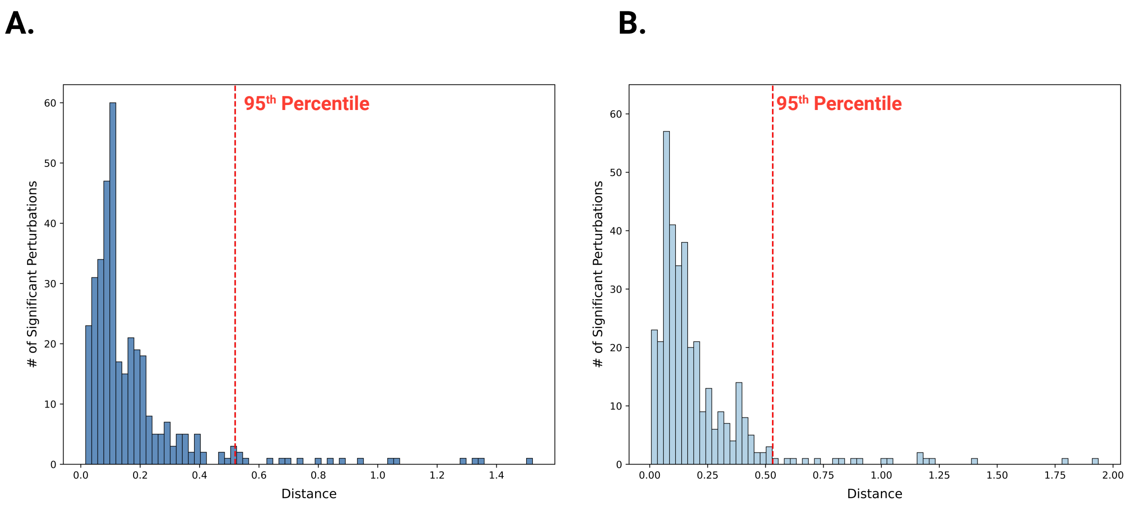


**Supplementary Figure S6: Distributions of distances for 2D projected simulations from the centroid. (A)** Distance distribution for $WT_{CAF}$. **(B)** Distance distribution for $WT_{CRC}$.

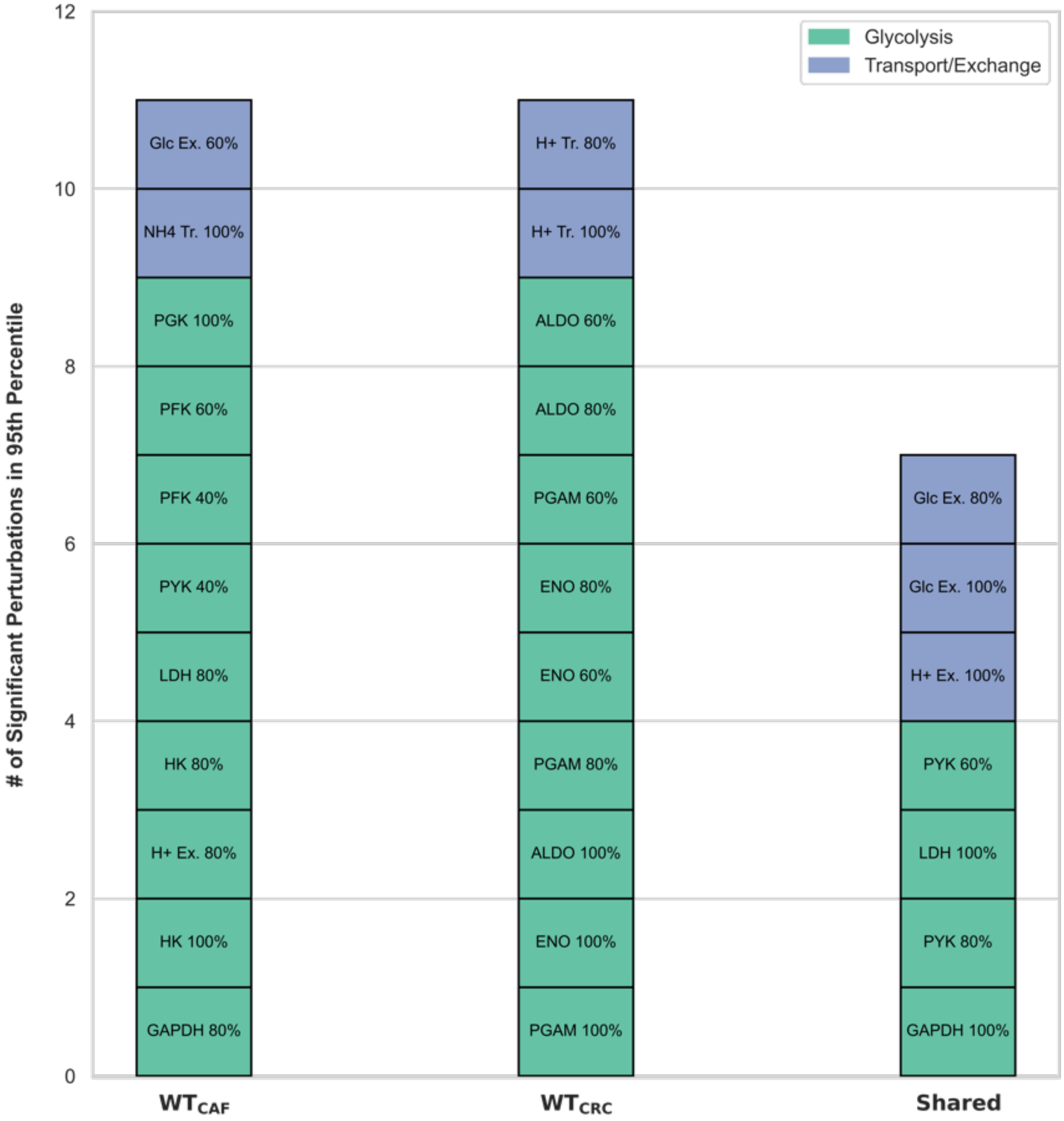


**Supplementary Figure S7: Pathway characterizations and comparisons of the significant perturbations between $WT_{CAF}$ and $WT_{CRC}$.**

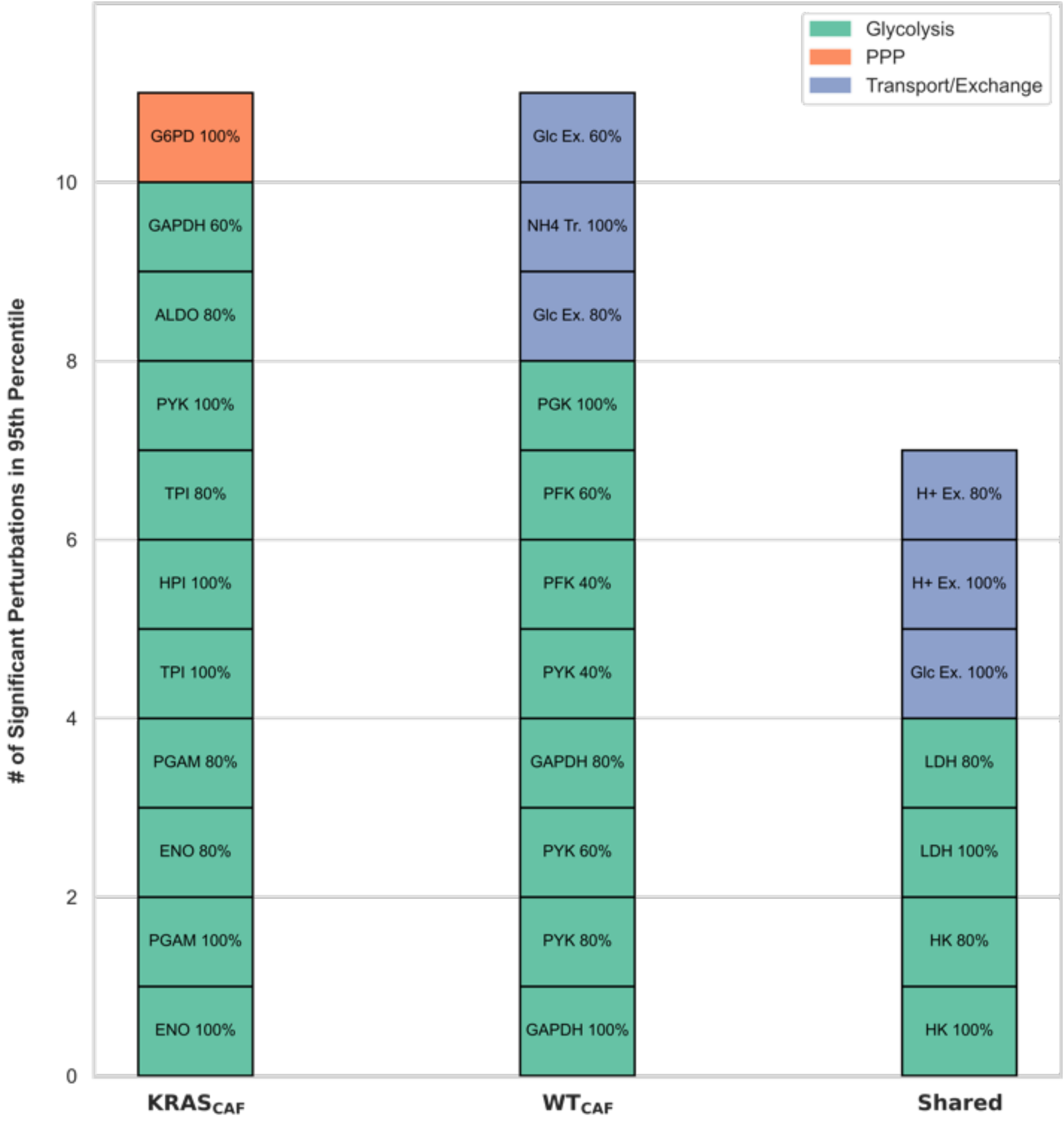


**Supplementary Figure S8: Pathway characterizations and comparisons of the significant perturbations between KRAS$_{CAF}$ and WT$_{CAF}$.**

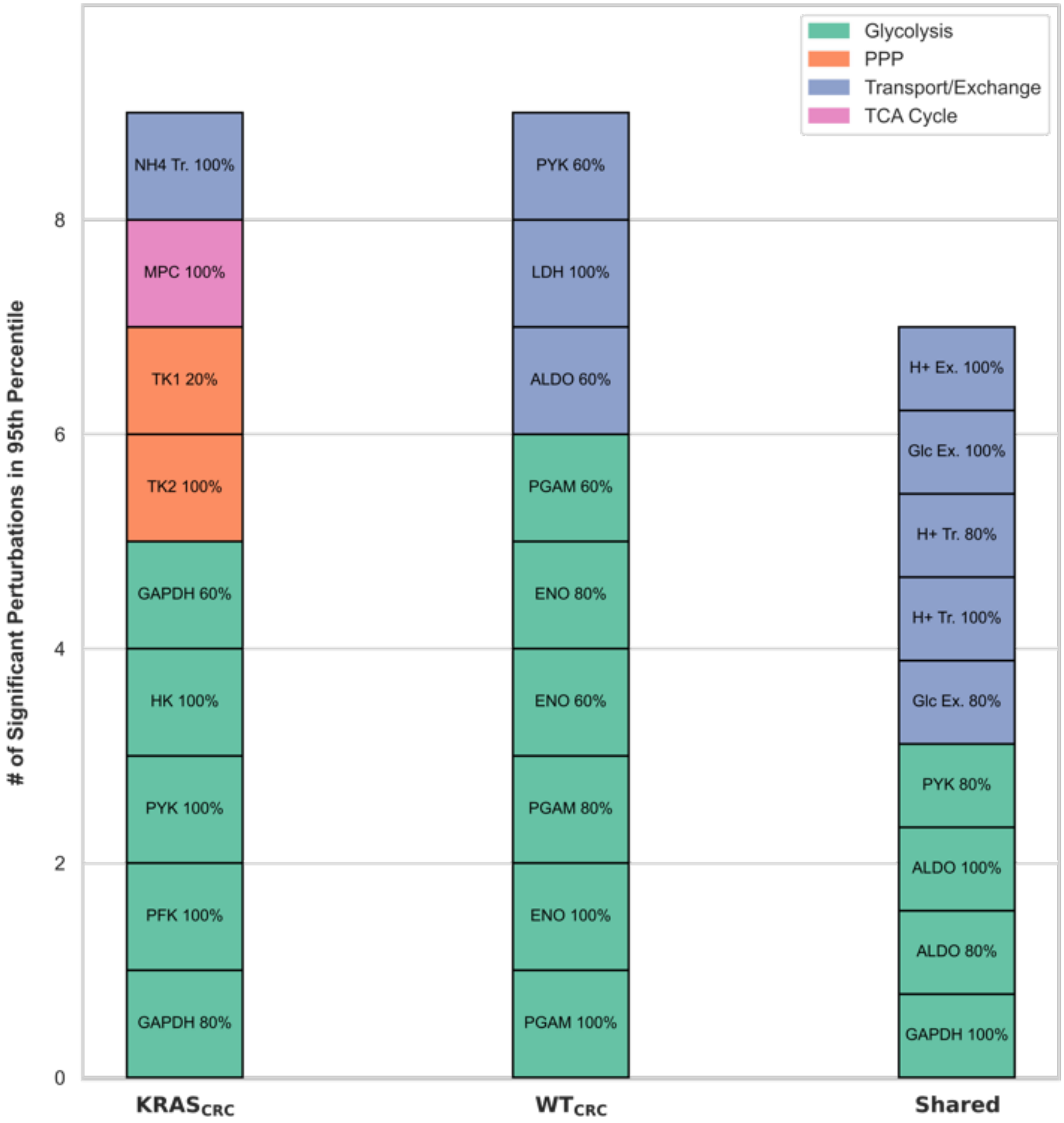


**Supplementary Figure S9: Pathway characterizations and comparisons of the significant perturbations between $KRAS_{CRC}$ and $WT_{CRC}$.**

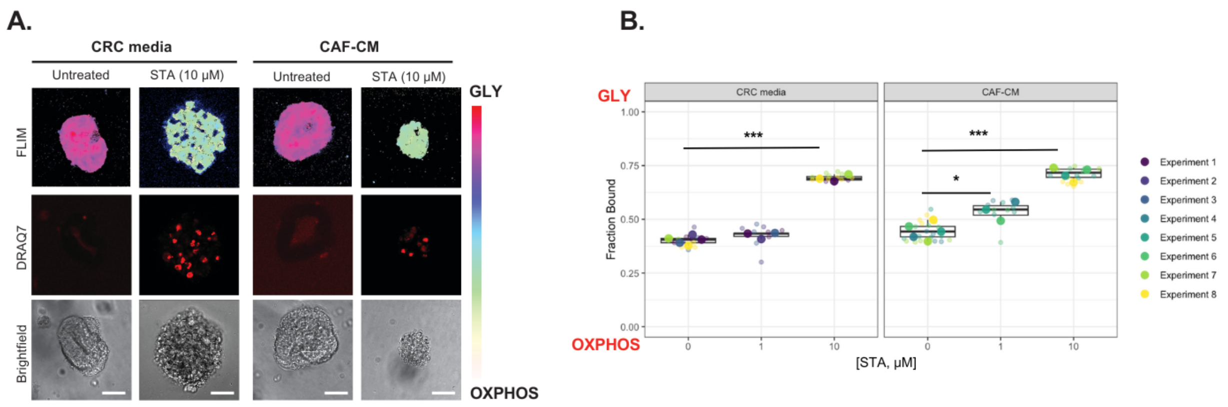


**Supplementary Figure S10: PDTOs treated with positive control staurosporine (STA). (A)** Representative FLIM images of PDTOs (000US) treated with STA in CRC media and CAF-CM and their **(B)** quantified metabolic signatures. For CAF-CM: n=4 (untreated); n=3 (1μM, 10μM). Scale bar is 20 μm.

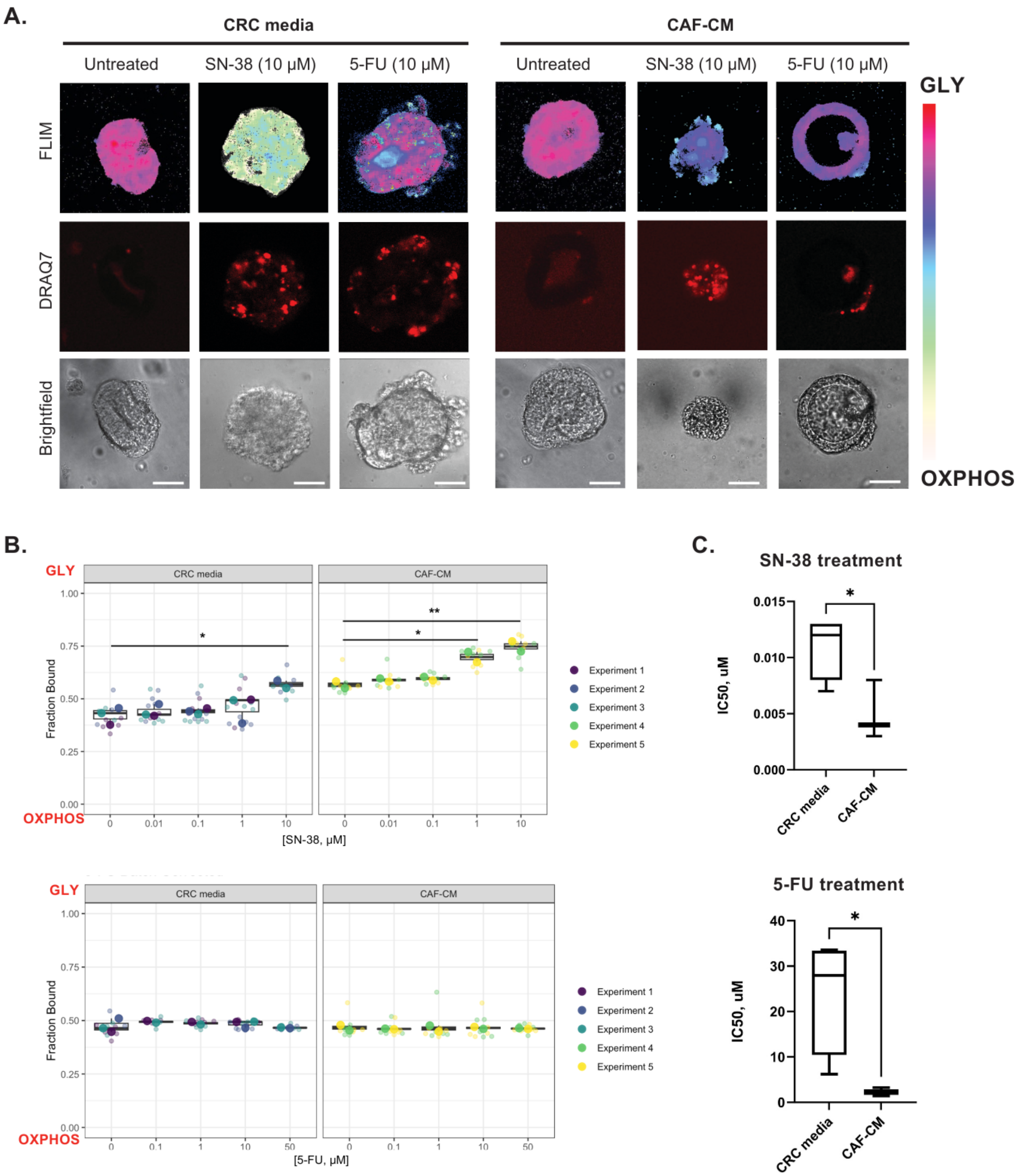


**Supplementary Figure S11: PDTOs treated with SN-38 and 5-FU. (A)** Representative FLIM images of PDTOs (000US) treated with SN-38 and 5-FU in CRC media and CAF-CM. Scale bar is 20 µm. **(B)** Quantified metabolic signature values for PTDOs treated with SN-38 or 5-FU in CRC media or CAF-CM. For SN-38 or 5-FU treated PDTOs in CAF-CM: n=2 for all conditions. For SN-38 in CRC media: n=3 (untreated, 0.01µM, 0.1µM, 1µM); n=2 (10µM). For 5-FU in CRC media: n= 3 (untreated, 10µM); n=2 (0.1µM, 1µM, and 50µM). **(C)** Calculated $IC_{50}$ values for dose response curves of SN-38 and 5-FU in CRC media or CAF-CM. *p<0.05, **p<0.01

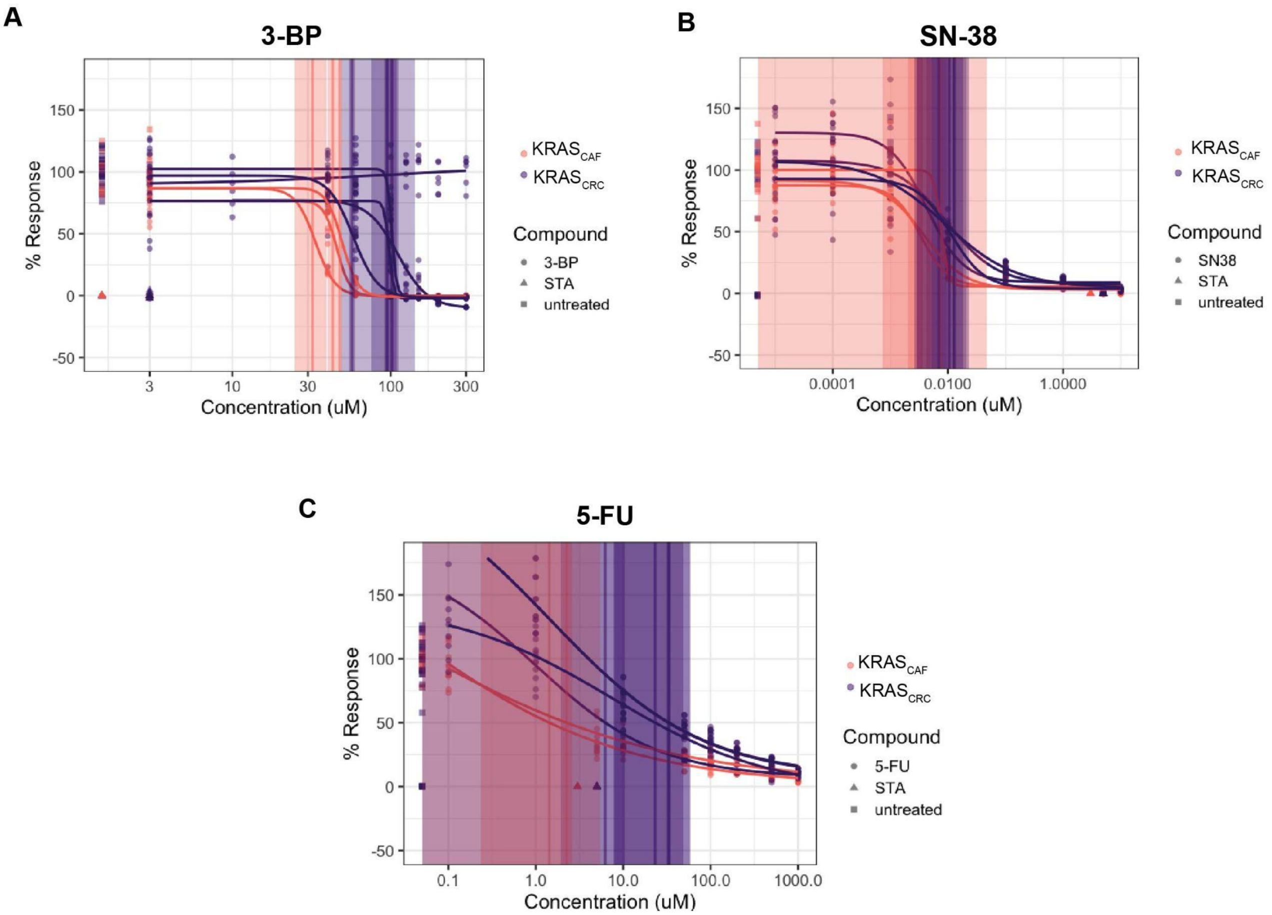


**Supplementary Figure S12 Full dose response curves of drug panel.** Dose response curves of PDTOs (000US) that passed QC metrics to calculate $IC_{50}$s for **(A)** 3-BP, **(B)** SN-38, and **(C)** 5-FU. In all plots, shaded regions correspond to 95% confidence intervals about the absolute $IC_{50}$s. QC metrics can be found in Supplementary Table S1.

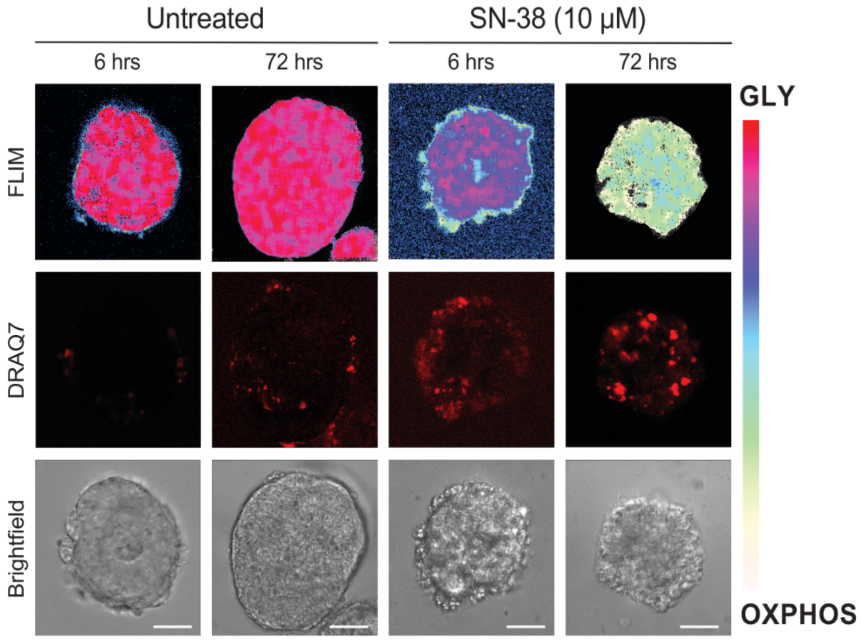


**Supplementary Figure S13: FLIM detects drug effects at earlier timepoints**. FLIM images of PDTOs after 6 and 72 hours of treatment. Positive DRAQ7 staining (red) indicates dead cells. Scale bar is 20 μm.

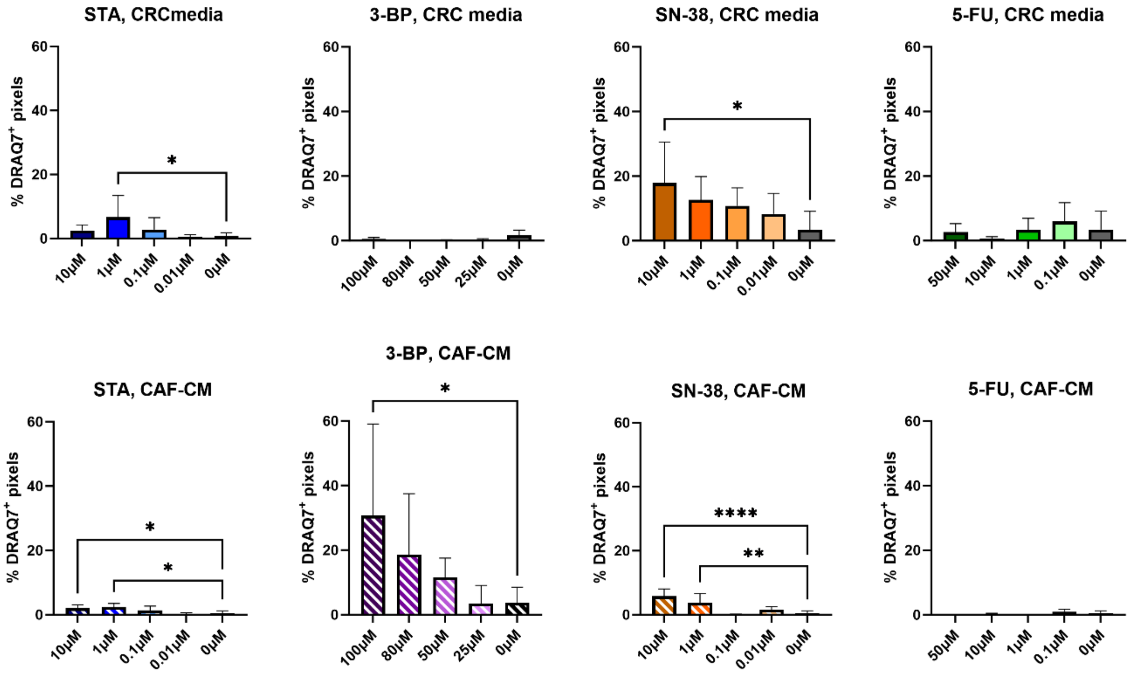


**Supplemental Figure S14: DRAQ7 quantification of images.** The percentage of DRAQ7$^+$ pixels were calculated by normalizing to the total number of pixels from only organoid from their respective brightfield images. * $p<0.05$

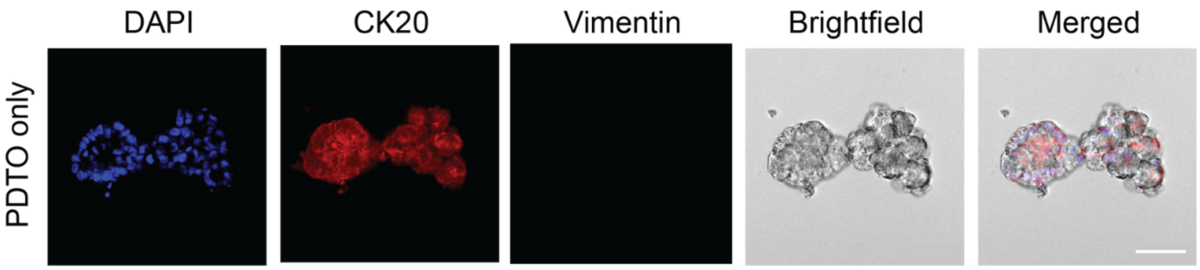


**Supplemental Figure S15: Immunostaining of PDTOs.** Organoids were stained for the nuclei (DAPI), colon tumor marker CK20, and stromal cell marker vimentin.

### Supplementary Tables

**Supplementary Table S1:** Utilized bounds for initial metabolite concentrations of $KRAS_{WT}$ and $KRAS_{MUT}$ CRC cells for upFBA, to predict flux distributions of the system. These ranges were taken from previously published data from human cervical cancer cells and breast cancer cell extracts. These ranges were increased and decreased by 20% to account for uncertainty (Wang et al., 2022).

| Metabolite | Lower (mM) | Upper (mM) |
|---|---|---|
| G6P [c] | 1.68 | 6 |
| FBP [c] | 0.16 | 1.08 |
| G3P [c] | 0.24 | 0.84 |
| PEP [c] | 0.24 | 0.84 |
| Lactate [c] | 6.4 | 19.2 |
| Gln [c] | 0.56 | 1.08 |
| Glu [m] | 0.88 | 1.56 |

**Supplementary Table S2:** Reported measurements of experimentally measured glucose uptake rates, lactate secretion rates, glutamine uptake rates, and cell proliferation rates. Glucose uptake, lactate secretion, and glutamine uptake rates between T = 0 hrs and T = 24 hours were measured via LC-MS metabolomics; biomass growth rates were measured via cell proliferation assays. The units for glucose uptake, lactate secretion, and glutamine uptake rates are mmM•$hr^{-1}$; the units for biomass growth rates are $hr^{-1}$. The numbers inside parentheses stand for standard deviations (Wang et al., 2022).

| | **Glucose ($mM•hr^{-1}$)** | **Lactate ($mM•hr^{-1}$)** | **Glutamine ($mM•hr^{-1}$)** | **Biomass ($hr^{-1}$)** |
|---|---|---|---|---|
| **$KRAS^{WT}$ (CRC media)** | 0.223 *(0.038)* | 0.283 *(0.016)* | 0.003 *(0.002)* | 0.035 *(0.001)* |
| **$KRAS^{MUT}$ (CRC media)** | 0.210 *(0.024)* | 0.234 *(0.004)* | 0.003 *(0.001)* | 0.034 *(0.003)* |
| **$KRAS^{WT}$ (CAF-cond. media)** | 0.573 *(0.087)* | 0.784 *(0.041)* | 0.052 *(0.025)* | 0.033 *(0.001)* |
| **$KRAS^{MUT}$ (CAF-cond. media)** | 0.927 *(0.135)* | 1.117 *(0.144)* | 0.105 *(0.049)* | 0.033 *(0.003)* |

**Supplementary Table S3: Sequencing results for PDTOs and respective tumor tissue (patient 000US).** Genetic mutations identified from DNA extracted from an intact patient tumor tissue sample or derived organoid cultures using Illumina's TruSight Oncology 500 next-generation sequencing platform.

| **Genetic Mutations Identified for CRC Patient** | |
|---|---|
| **Mutations Present In Both Tumor Tissue & PDOs** | |
| **Gene Symbol** | **Protein Change** |
| *KRAS* | p.Gly12Ala |
| *TP53* | p.Gly245Ser |
| *SMAD4* | p.Arg361Cys |
| *ERBB4* | p.Arg103Cys |
| *SH2B3* | p.Ser213Arg |
| *CIC* | p.Arg415Trp |
| *MCL1* | p.Val76Ile |
| *PTPRD* | p.Gln67His |
| *GATA6* | p.Pro144Arg |
| *INPP4A* | p.Lys412Arg |
| **Mutations Present In Tumor Tissue Only** | |
| *MAP3K1* | p.Arg315Lys |
| *AXL* | p.Pro771His |
| *PBRM1* | p.Pro1429Ser |
| *AMER1* | p.Glu762Ter |
| *NCOR1* | p.Thr525_Lys527del |
| **Mutations Present In PDTOs Only** | |
| *FLT3* | p.Arg773GlyfsTer8 |
| *EP300* | p.Leu1160Phe |
| *ZFHX3* | p.Gln3380del |
| *KAT6A* | p.Ser396Asn |
| *STAG1* | p.Glu433Ter |

**Supplementary Table S4:** Summary of dose-response curve QC metrics.

| Media | Drug | Experiment | IC50 (uM) | RSE | Average IC50 | Average RSE |
|---|---|---|---|---|---|---|
| CRC media | SN38 | Run 1 | 0.007 | 13.775 | | |
| | | Run 2 | 0.013 | 14.196 | | |
| | | Run 3 | 0.013 | 15.911 | | |
| | | Run 4 | 0.011 | 22.781 | 0.011 | 16.666 |
| | 5FU | Run 1 | 33.570 | 11.990 | | |
| | | Run 2 | 6.230 | 12.910 | | |
| | | Run 3 | 23.160 | 12.930 | | |
| | | Run 4 | 32.770 | 12.090 | 23.933 | 12.480 |
| | 3BP | Run 1 | 93.410 | 16.970 | | |
| | | Run 2 | 101.980 | 8.240 | | |
| | | Run 3 | 95.420 | 18.070 | | |
| | | Run 4 | 57.040 | 6.100 | 86.963 | 12.345 |
| CAF-CM | SN38 | Run 1 | 0.004 | 10.019 | | |
| | | Run 2 | 0.008 | 10.900 | | |
| | | Run 3 | 0.003 | 14.855 | 0.005 | 11.925 |
| | 5FU | Run 1 | 1.430 | 6.919 | | |
| | | Run 2 | 2.250 | 9.130 | | |
| | | Run 3 | 3.240 | 4.570 | 2.307 | 6.873 |
| | 3BP | Run 1 | 32.050 | 4.630 | | |
| | | Run 2 | 43.090 | 8.900 | | |
| | | Run 3 | 47.640 | 4.190 | 40.927 | 5.907 |

### Description of distances in 74-dimensional space

In the original high-dimensional, 74D space, the distances from the simulation vectors to a base (unperturbed) vector form a left-tailed distribution in which most of the distances are large and similar in number. As the number of dimensions increases, the volume of the space increases exponentially such that the available data become sparse. This sparsity is why distances in high-dimensional spaces are often larger and more varied than in low-dimensional spaces. Furthermore, in higher-dimensional spaces, the contrast between near and far points can be exaggerated, leading to a distribution with more large values.

Conversely, after reducing the dimensionality to 2D, the distances from the data points to the cluster centers yield a right-tailed distribution. This means that the majority of data points are nearer to the centers, with only a few outliers. The significant change in the distribution shape from 74D to 2D is attributed to the biases inherent in the high-dimensional space, where all pairwise distances are weighted equally. In 2D, representation learning allows for differential weighting, giving more or less importance to the distances based on the similarity or dissimilarity of each data point, under specific conditions such as partial knockdowns. The dimensionality reduction process tries to maintain the most significant structures, usually by preserving the local neighborhoods, which can lead to a more clustered appearance with fewer "far apart" distances.

Thus, dimensionality reduction via representation learning refines the significance of distances between data points, enabling us to use these distances as reliable indicators of the similarities or differences between simulations in a reduced dimensionality space.
